# Imprinted SARS-CoV-2 humoral immunity induces convergent Omicron RBD evolution

**DOI:** 10.1101/2022.09.15.507787

**Authors:** Yunlong Cao, Fanchong Jian, Jing Wang, Yuanling Yu, Weiliang Song, Ayijiang Yisimayi, Jing Wang, Ran An, Xiaosu Chen, Na Zhang, Yao Wang, Peng Wang, Lijuan Zhao, Haiyan Sun, Lingling Yu, Sijie Yang, Xiao Niu, Tianhe Xiao, Qingqing Gu, Fei Shao, Xiaohua Hao, Yanli Xu, Ronghua Jin, Zhongyang Shen, Youchun Wang, Xiaoliang Sunney Xie

## Abstract

Continuous evolution of Omicron has led to a rapid and simultaneous emergence of numerous variants that display growth advantages over BA. 5. Despite their divergent evolutionary courses, mutations on their receptor-binding domain (RBD) converge on several hotspots. The driving force and destination of such convergent evolution and its impact on humoral immunity remain unclear. Here, we demonstrate that these convergent mutations can cause striking evasion of neutralizing antibody (NAb) drugs and convalescent plasma, including those from BA.5 breakthrough infection, while maintaining sufficient ACE2 binding capability. BQ.1.1.10, BA.4.6.3, XBB, and CH. 1.1 are the most antibody-evasive strain tested, even exceeding SARS-CoV-1 level. To delineate the origin of the convergent evolution, we determined the escape mutation profiles and neutralization activity of monoclonal antibodies (mAbs) isolated from BA.2 and BA.5 breakthrough-infection convalescents. Importantly, due to humoral immune imprinting, BA.2 and especially BA.5 breakthrough infection caused significant reductions in the epitope diversity of NAbs and increased proportion of non-neutralizing mAbs, which in turn concentrated humoral immune pressure and promoted convergent evolution. Moreover, we showed that the convergent RBD mutations could be accurately inferred by integrated deep mutational scanning (DMS) profiles, and the evolution trends of BA.2.75/BA.5 subvariants could be well-simulated through constructed convergent pseudovirus mutants. Together, our results suggest current herd immunity and BA.5 vaccine boosters may not provide good protection against infection. Broad-spectrum SARS-CoV-2 vaccines and NAb drugs development should be highly prioritized, and the constructed mutants could help to examine their effectiveness in advance.

## Main

SARS-CoV-2 Omicron BA.1, BA.2, and BA.5 have demonstrated strong neutralization evasion capability, posing severe challenges to the efficacy of existing humoral immunity established through vaccination and infection ^1–14^. Nevertheless, Omicron is continuously evolving, leading to various new subvariants, including BA.2.75, BA.4.6, and BF.7 ^15–21^. Importantly, a high proportion of these emerging variants display significant growth advantages over BA.5, such as BA.2.3.20, BA.2.75.2, BQ.1.1, and especially XBB, a recombinant of BJ.1 and BM.1.1.1 (Fig. 1) ^22^. Such rapid and simultaneous emergence of multiple variants with enormous growth advantages is unprecedented. Notably, although these derivative subvariants appear to diverge along the evolutionary course, the mutations they carry on the receptor-binding domain (RBD) converge on the same sites, including R346, K356, K444, V445, G446, N450, L452, N460, F486, F490, R493, and S494 (Fig. 1). Most mutations on these residues are known to be antibody-evasive, as revealed by deep mutational scanning (DMS) ^1,2,23–25^. It’s crucial to examine the impact of these convergent mutations on antibody-escaping capability, receptor binding affinity, and the efficacy of vaccines and antibody therapeutics. It’s also important to investigate the driving force behind this suddenly accelerated emergence of RBD mutations, what such mutational convergence would lead to, and how we can prepare for such convergent RBD evolution.

**Fig. 1.**
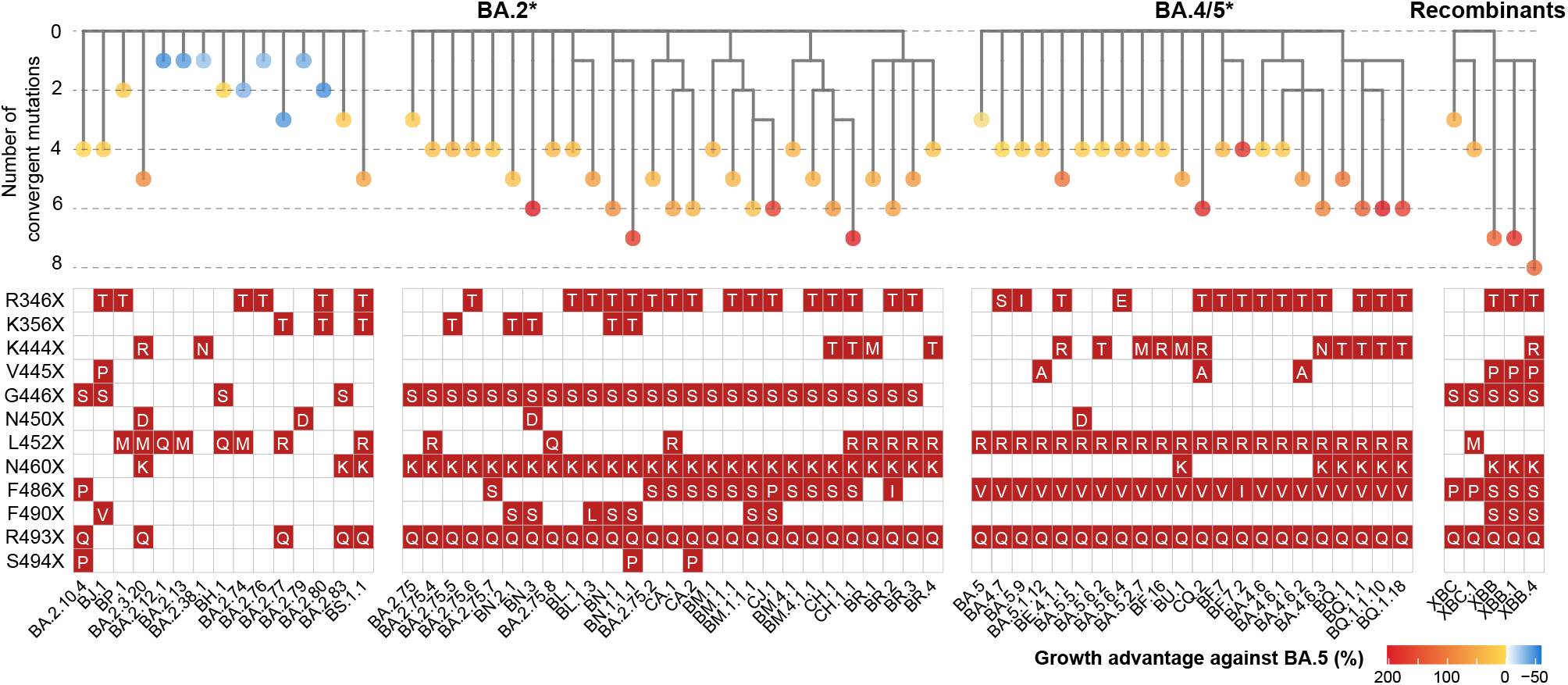
Convergent evolution of Omicron RBD with growth advantage over BA.5. Phylogenetic tree of featured Omicron subvariants carrying convergent mutations. Their relative growth advantage values calculated using the CoV-Spectrum website are indicated as color scale. Specific convergent mutations carried by each lineage are labeled.

### NAb evasion caused by convergent variants

First, we tested the antibody evasion capability of these convergent variants. We constructed the VSV-based spike-pseudotyped virus of Omicron BA.2, BA.2.75, and BA.4/5 sublineages carrying those convergent mutations and examined the neutralizing activities of therapeutic neutralizing antibodies (NAbs) against them (Fig. 2a and Extended Data Fig. 1a) ^26–28^. In total, pseudoviruses of 50 convergent variants were constructed and tested. COV2-2196+COV2-2130 (Evusheld) ^26,29^ is vulnerable to F486, R346, and K444-G446 mutations, evaded or highly impaired by BJ.1 (R346T), XBB (R346T+V445P+F486S), BA.2.75.2/CA.1/BM.1.1/BM.1.1.1/CH.1.1 (R346T+F486S), CJ.1 (R346T+F486P), BR.2 (R346T+F486I), BA.4.6.1 (R346T+F486V), BA.5.6.2/BQ.1 (K444T+F486V), BU.1 (K444M+F486V), and BQ.1.1 (R346T+K444T+F486V). LY-CoV1404 (Bebtelovimab) remains potent against BF.16 (K444R) and BA.5.5.1 (N450D) and shows reduced potency against BA.5.1.12 (V445A) ^27^ (Extended Data Fig. 1a). However, LY-CoV1404 was escaped by BJ.1, XBB, BR.1, CH.1.1, BA.4.6.3 and BQ.1.1 while exhibiting strongly reduced activity against BA.2.38.1, BA.5.6.2, and BQ.1 due to K444N/T mutations and the combination of K444M/G446S or V445P/G446S ^27^. SA55+SA58 is a pair of broad NAbs isolated from vaccinated SARS convalescents that target non-competing conserved epitopes ^2,28^. SA58 is weak to G339H and R346T mutations and showed reduced neutralization efficacy against BJ. 1/XBB and BA.2.75 sublineages. SA55 is the only NAb demonstrating high potency against all tested Omicron subvariants. Among the tested variants, XBB and BQ.1.1 exhibited the strongest resistance to therapeutic mAbs and cocktails (Fig. 2a). Since the SA55+SA58 cocktail is still in preclinical development, the efficacy of available antibody drugs, including the BA.2.75/BA.5-effective Evusheld and Bebtelovimab, are extensively affected by the emerging subvariants with convergent mutations.

**Fig. 2.**
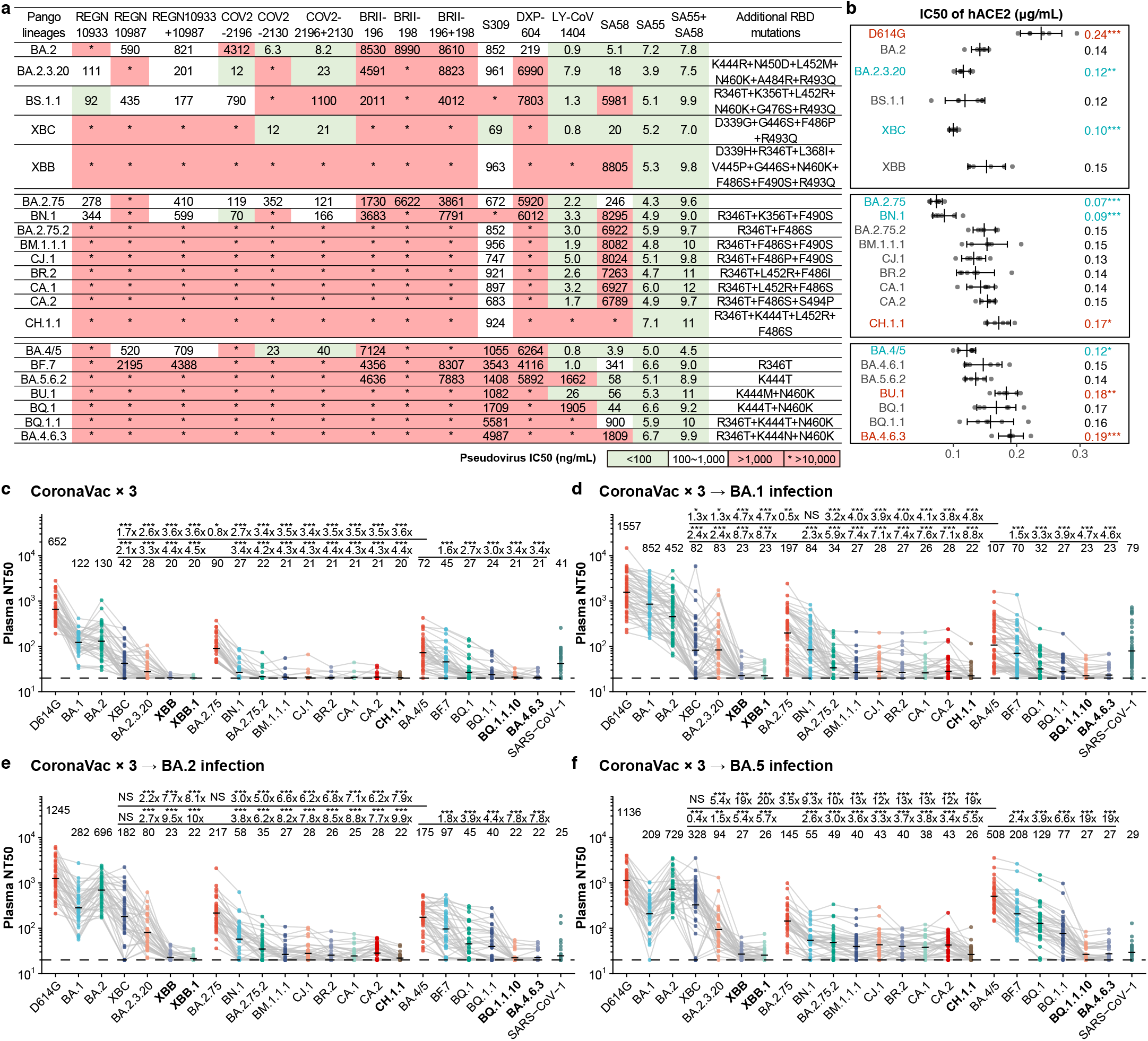
Convergent Omicron subvariants induce striking NAb evasion. **a**, IC50 of therapeutic NAbs against VSV-based pseudoviruses with spike glycoproteins of emerging SARS-CoV-2 BA.2/BA.5/BA.2.75 convergent subvariants. green, IC50 ≤ 100ng/mL; white, 100ng/mL < IC50 < 1,000ng/mL; red, IC50 ≥ 1,000ng/mL; *, IC50 ≥ 10,000ng/mL. **b**, Relative hACE2-binding capability measured by IC50 of hACE2 against pseudoviruses of variants. Error bars indicate mean±s.d. P-values were calculated using twotailed Student’s *t*-test. *, p < 0.05; **, p < 0.01; ***, p < 0.001. No label on variants with p > 0.05. Variants with significantly stronger binding are colored blue, while those with weaker binding are colored red. **c-f**, Pseudovirus-neutralizing titers against SARS-CoV-2 D614G and Omicron subvariants of plasma from vaccinated individuals or convalescents of breakthrough infection. **c**, Individuals who had received 3 doses of CoronaVac (n = 40). **d**, Convalescents who had been infected with BA.1 after receiving 3 doses of CoronaVac (n = 50). **e**, Convalescents who had been infected with BA.2 after receiving 3 doses of CoronaVac (n = 39). **f**, Convalescents who had been infected with BA.5 after receiving 3 doses of CoronaVac (n = 36). The geometric mean titers are labeled. Statistical tests are performed using two-tailed Wilcoxon signed-rank tests of paired samples. *, p < 0.05; **, p < 0.01; ***, p < 0.001; NS, not significant, p > 0.05. NT50 against BA.2 and BA.2.75-derived subvariants are compared to both that against BA.4/5 (the first line) and BA.2.75 (the second line); while BA.4/5-derived subvariants are only compared with BA.4/5.

Sufficient ACE2-binding affinity is essential for SARS-CoV-2 transmission. Thus, we examined the relative hACE2 binding capability of these variants by evaluating hACE2 inhibitory efficiency against the pseudoviruses. Higher inhibitory efficiency of soluble hACE2 against pseudoviruses indicates higher ACE2-binding capability ^17^. Overall, these convergent variants all demonstrate sufficient ACE2-binding efficiency, at least higher than that of D614G, including the most antibody-evasive XBB, BQ.1.1, and CH.1.1 (Fig. 2b and Extended Data Fig. 1b). Specifically, R493Q reversion increases the inhibitory efficiency of hACE2, which is consistent with previous reports ^3,17,19^. K417T shows a moderate increase in the inhibitory efficiency of hACE2. In contrast, F486S, K444M, and K444N have a clear negative impact on inhibitory efficiency, while K444T and F486P do not cause significant impairment of ACE2 binding. These observations are also in line with previous DMS results ^30^.

Most importantly, we investigated how these variants escape the neutralization of plasma samples from individuals with various immune histories. We recruited cohorts of individuals who received 3 doses of CoronaVac ^31^ with or without breakthrough infection by BA.1, BA.2, or BA.5. Convalescent plasma samples were collected on average around 4 weeks after hospital discharge (Supplementary Table 1). Plasma from CoronaVac vaccinees was obtained 4 weeks after the third dose. A significant reduction in the NT50 against most tested BA.2, BA.2.75, or BA.5 subvariants has been observed, compared to that against corresponding ancestral BA.2, BA.2.75, or BA.5, respectively (Fig. 2c-f and Extended Data Fig. 2a-d).

Specifically, BA.2.3.20 and BA.2.75.2 are significantly more immune evasive than BA.5 (Fig. 2c-f), explaining their high growth advantage. Nevertheless, multiple convergent variants showed even stronger antibody evasion capability, including (BM.1.1+F490S), CJ. 1 (BM.1.1.1+S486P), CA.1 (BA.2.75.2+L452R+T604I), CA.2 (BA.2.75.2+S494P), CH.1 (BA.2.75+R346T+K444T+F486S), CH.1.1 (CH.1+L452R) in the BA.2.75 sublineages, and BQ. 1.1 (BA.5+R346T+K444T+N460K), BQ. 1.1.10 (BQ.1.1+Y144del), and BA.4.6.3 (BA.4.6+K444N+N460K+Y144del) in the BA.4/5 sublineages. Strikingly, the BJ.1/BM.1.1.1 recombinant strain XBB and XBB.1 (XBB+G252V) are among the most humoral immune evasive strains tested, comparable to that of CH.1.1, BQ.1.1.10 and BA.4.6.3. Importantly, BA.5 breakthrough infection yields higher plasma NT50 against BA.5 sublineages, including BQ. 1.1; however, plasma from BA.5 breakthrough infection neutralize poorly against XBB, CH. 1.1, BQ. 1.1.10 and BA.4.6.3, suggesting that the NTD mutations they carry are extremely effective at evading NAbs elicited by BA.5 breakthrough infection (Fig. 2f). Notably, the strongest immune-evasive convergent variants have displayed even lower NT50 than SARS-CoV-1, suggesting immense antigenic drift and potential serotype conversion.

### Immune imprinting induces convergent evolution

It is crucial to investigate the origin of such accelerated RBD convergent evolution. Therefore, we characterized the antibody repertoires induced by Omicron BA.2 and BA.5 breakthrough infection, which is the dominant immune background of current global herd immunity. Following the strategy described in our previous report using pooled PBMC from BA.1 breakthrough infection ^2^, we enriched antigen-specific memory B cells by fluorescence-activated cell sorting (FACS) for individuals who had recovered from BA.2 and BA.5 breakthrough infection (Supplementary Table 1). RBD-binding CD27^+^/IgM^-^/IgD^-^ cells were subjected to single-cell V(D)J sequencing (scVDJ-seq) to determine the BCR sequences (Extended Data Fig. 3a-b).

Similar to that reported in BA.1 breakthrough infection, immune imprinting, or so-called “original antigenic sin”, is also observed in BA.2 and BA.5 breakthrough infection ^2,32–35^. Postvaccination infection with BA.2 and BA.5 mainly recalls cross-reactive memory B cells elicited by wildtype-based vaccine, but rarely produces BA.2/BA.5 specific B cells, similar to BA. 1 breakthrough infection (Fig. 3a-b). This is in marked contrast to Omicron infection without previous vaccination (Fig. 3c and Extended Data Fig. 3c). The RBD-targeting antibody sequences determined by scVDJ-seq are then expressed *in vitro* as human IgG1 monoclonal antibodies (mAbs). As expected, only a small proportion of the expressed mAbs specifically bind to BA.2/BA.5 RBD and are not cross-reactive to WT RBD, determined by enzyme-linked immunosorbent assay (ELISA), concordant with the FACS results (Fig. 3d). Importantly, crossreactive mAbs exhibit significantly higher somatic hypermutation (SHM), indicating that these antibodies are more affinity-matured and are indeed most likely recalled from previously vaccination-induced memory (Fig. 3d).

**Fig. 3.**
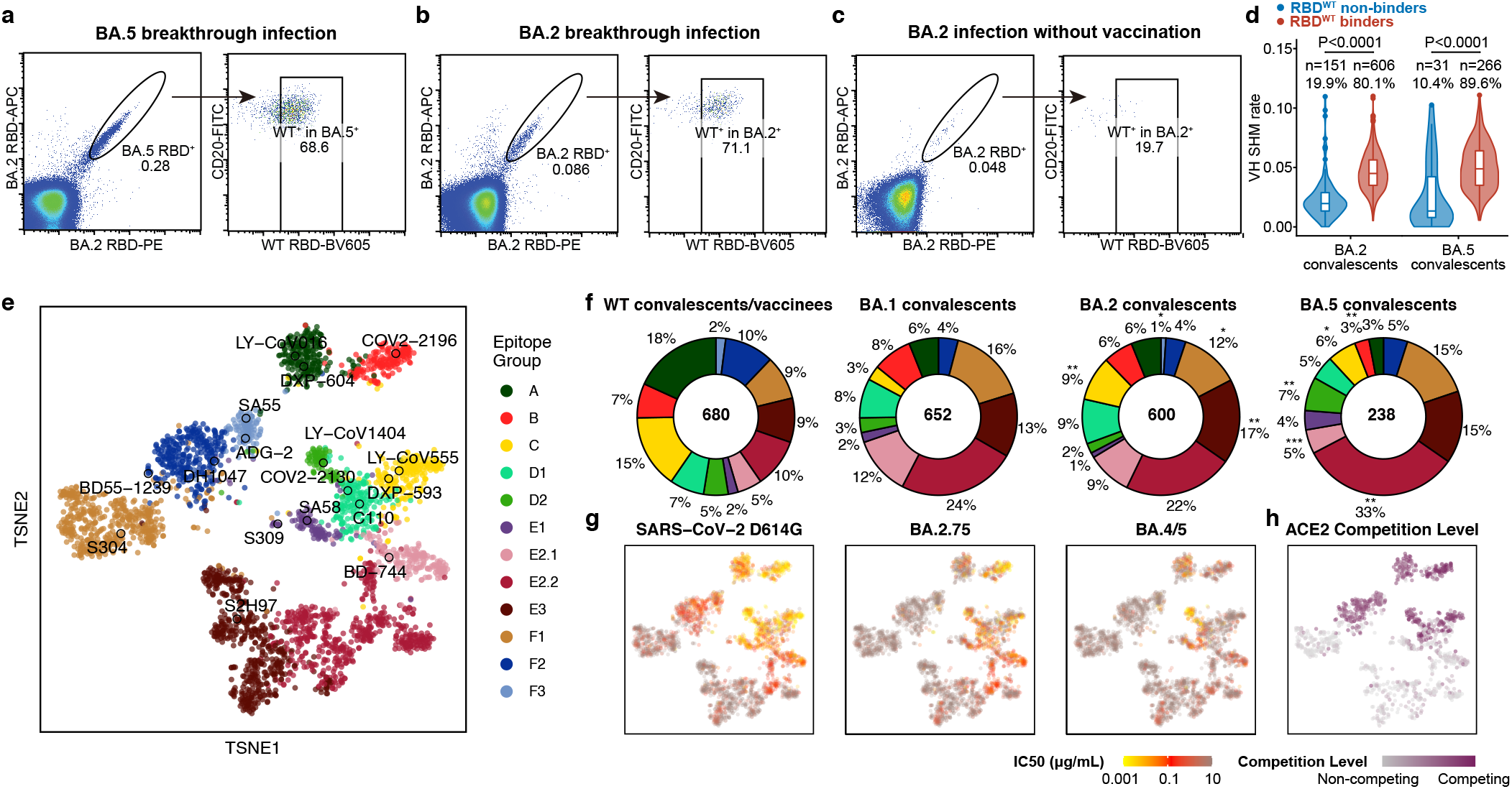
Epitope characterization of mAbs elicited by Omicron breakthrough convalescents. **a-c**, FACS analysis of pooled memory B cells (IgM^-^, IgD^-^/CD27^+^) from Omicron convalescents. **a**, BA.5 breakthrough infection; **b**, BA.2 breakthrough infection; **c**, BA.2 convalescents without vaccination. **d**, The heavy chain variable domain SHM rate of mAbs from BA.2 and BA.5 breakthrough infection convalescents. Binding specificity is determined by ELISA. Statistical tests are determined using two-tailed Wilcoxon signed-rank tests. Boxes display the 25th percentile, median and 75th percentile, and violin plots show kernel density estimation curves of the distribution. **e**, t-SNE and clustering of SARS-CoV-2 wildtype RBD-binding antibodies based on DMS profiles of 3051 antibodies. **f**, Epitope distribution of mAbs from wildtype convalescents and post-vaccination BA.1/BA.2/BA.5 infection convalescents. Twotailed binomial tests were used to compare the proportion of each epitope group from BA.2 and BA.5 convalescents with that from BA.1. *, p < 0.05; **, p < 0.01; ***, p < 0.001; No label for p > 0.05. **g**, Neutralizing activity projection of mAbs against SARS-CoV-2 D614G (n=3046), BA.2.75 (n=3046), and BA.4/5 (n=3046), respectively. **h**, ACE2 competition level projection of mAbs determined by competition ELISA (n=1317).

Next, we determined the escape mutation profiles of these antibodies by high-throughput DMS and measured their neutralizing activities against SARS-CoV-2 D614G, BA.2, BA.5, BA.2.75, BQ.1.1 and XBB (Fig. 3e, 3g and Extended Data Fig. 4a-b). Previously, we reported the DMS profiles and the epitope distribution of antibodies isolated from WT vaccinated/infected individuals, SARS-CoV-2 vaccinated SARS convalescents, and BA.1 convalescents, which could be classified into 12 epitope groups ^2^. Among them, mAbs in groups A, B, C, D1, D2, F2, and F3 compete with ACE2 and exhibit neutralizing activity (Fig. 3h, Extended Data Fig. 5a-d and 6a-d); while mAbs in groups E1, E2.1, E2.2, E3, and F1 do not compete with ACE2 (Fig. 3h, Extended Data Fig. 7a-c). Antibodies in groups E2.2, E3, and F1 exhibit low or no neutralizing capability (Extended Data Fig. 4b, 7d). To integrate the previous dataset with DMS profiles of the new mAbs isolated from BA.2 and BA.5 convalescents, we co-embedded all antibodies using multidimensional scaling (MDS) based on their DMS profiles, followed by *t*-distributed stochastic neighbor embedding (*t*-SNE) for visualization, and used KNN-based classification to determine the epitope groups of new mAbs (Fig. 3e). This results in a dataset containing the DMS profiles of 3051 SARS-CoV-2 WT RBD-targeting mAbs in total (Supplementary Table 2). The epitope distribution of mAbs from BA.2 breakthrough infection is generally similar to those elicited by BA.1, except for the increased proportion of mAbs in group C (Fig. 3f). However, BA.5-elicited mAbs showed a more distinct distribution compared to BA.1, with a significantly increased proportion of mAbs in group D2 and E2.2, and decreased ratio of antibodies in groups B and E2.1. The main reason is that the F486 and L452 mutations carried by BA.5 make these cross-reactive memory B cells unable to be activated and recalled (Fig. 3f, Extended Data Fig. 5b, 6a and 7a). Remarkably, antibody repertoires induced by all Omicron breakthrough infections are distinct from those stimulated by WT. Compared to WT infection or vaccination, BA.1, BA.2, and BA.5 breakthrough infection mainly elicit mAbs of group E2.2, E3, and F1, which do not compete with ACE2 and demonstrate weak neutralizing activity, while WT-elicited antibodies enrich mAbs of groups A, B, and C which compete with ACE2 and exhibit strong neutralization potency (Fig. 3f-h). Strikingly, the combined proportion of E2.2, E3, and F1 antibodies rose from 29% in WT convalescents/vaccinees, 53% in BA.1 convalescents, 51% in BA.2 convalescents, to 63% in BA.5 convalescents (Fig. 3f). Overall, the proportion and diversity of neutralizing antibody epitopes are reduced in Omicron breakthrough infection, especially in BA.5 breakthrough infection.

To better delineate the impact of immune imprinting and consequent reduction of NAb epitope diversity on the RBD evolutionary pressure, we aggregated the DMS profiles of large collections of mAbs to estimate the impact of mutations on the efficacy of humoral immunity, as inspired by previous works (Supplementary Table 2) ^36^. It is essential to incorporate the effects of ACE2 binding, RBD expression, neutralizing activity of mAbs, and codon usage constraint with the escape profiles to estimate the SARS-CoV-2 evolution trend on the RBD. In brief, each mutation on the RBD would have an impact on each mAb in the set, which is quantified by the escape scores determined by DMS and weighted by its IC50 against the evolving strain. For each residue, only those amino acids that are accessible by one nucleotide mutation are included. Impacts on ACE2-binding capability (as measured by pseudovirus inhibitory efficiency) and RBD expression of each mutation are also considered in the analyses, using data determined by DMS in previous reports ^30,37,38^. Finally, the estimated relative preference of each mutation is calculated using the sum of weighted escape scores of all mAbs in the specific set.

The reduced NAb epitope diversity caused by imprinted humoral response could be strikingly shown by the estimated mutation preference spectrum (Fig. 4a). Diversified escaping-score peaks, which also represent immune pressure, could be observed when using BA.2-elicited antibodies, while only two major peaks could be identified, R346T/S and K444E/Q/N/T/M, when using BA.5-elicited antibodies (Fig. 4a). Interestingly, these two hotspots are the most frequently mutated sites in continuously evolving BA.4/5 subvariants, and convergently occurred in multiple lineages (Fig. 1). Similar analyses for WT and BA.1 also demonstrated diversified peaks; thus, the concentrated immune pressure strikingly reflects the reduced diversity of NAbs elicited by BA.5 breakthrough infection due to immune imprinting, and these concentrated preferred mutations highly overlapped with convergent hotspots observed in the real world (Extended Data Fig. 8a-b). Together, our results indicate that due to immune imprinting, BA.5 breakthrough infection caused significant reductions of NAb epitope diversity and increased proportion of non-neutralizing mAbs, which in turn concentrated immune pressure and promoted the convergent RBD evolution.

**Fig. 4.**
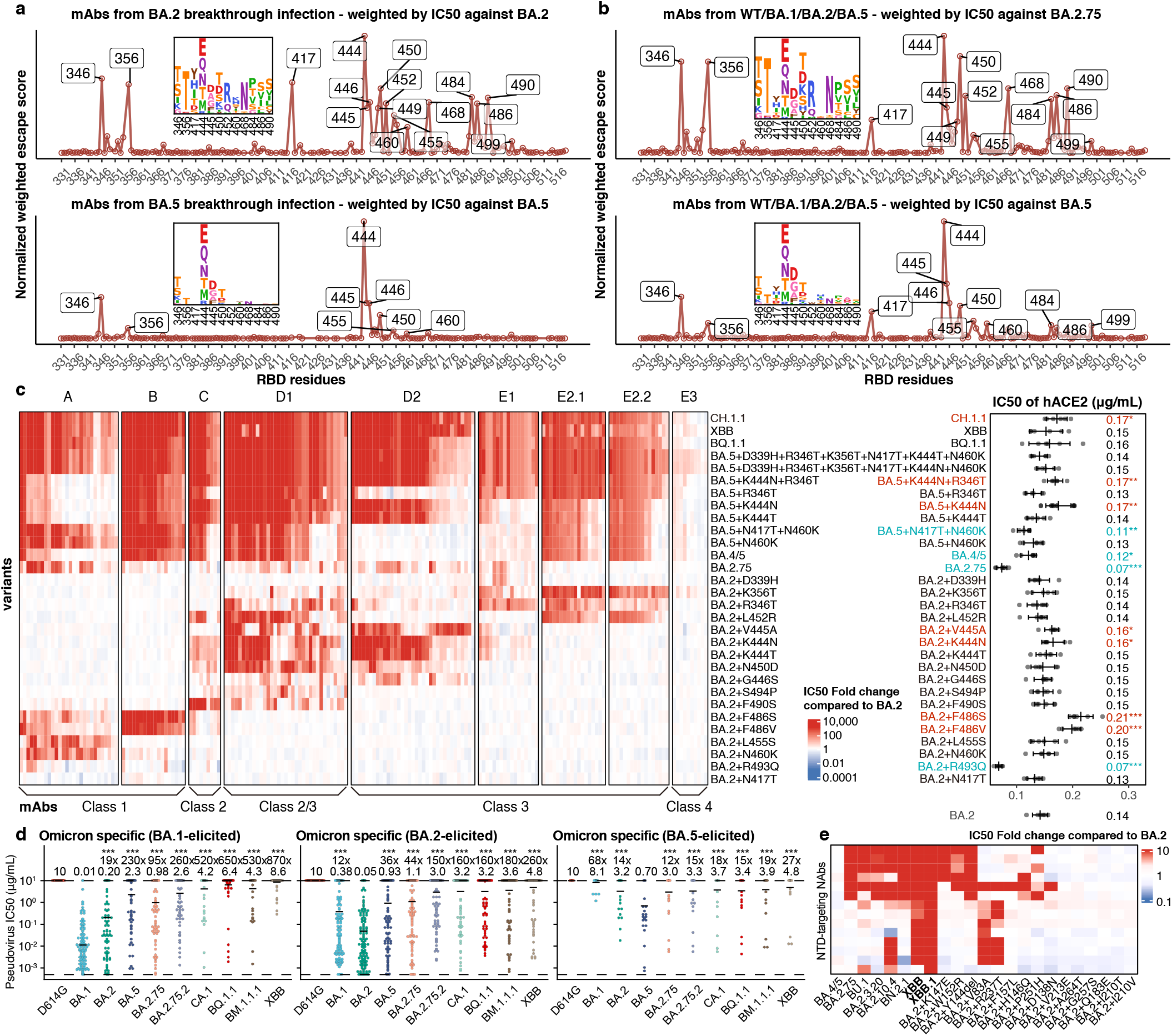
Immune imprinting promotes convergent evolution of NAb-evasive mutations. **a-b**, Normalized average escape scores weighted by IC50 against **a**, BA.2 and BA.5 using DMS profiles of NAbs from corresponding convalescents. **b**, BA.2.75 and BA.5 using DMS profiles of all NAbs except those from SARS convalescents. **c**, IC50 of representative potent BA.2-neutralizing antibodies in epitope group against emerging and constructed Omicron subvariants pseudovirus with escape mutations, in addition to IC50 of hACE2 against these variants. Classes of the NAbs as defined by Barnes et al. ^50^ are also annotated below this map. Error bars indicate mean±s.d. P-values were calculated using two-tailed Student’s t-test. *, p < 0.05; **, p < 0.01; ***, p < 0.001. No label on variants with p > 0.05. Variants with significantly stronger binding are colored blue, while those with weaker binding are colored red. **d**, IC50 against featured subvariants of RBD-targeting Omicron-specific NAbs from BA.1 (N=108), BA.2 (N=92), and BA.5 (N=17) breakthrough convalescents. The geometric mean IC50s are labeled, and error bars indicate the geometric standard deviation. P-values are calculated using twotailed Wilcoxon signed-rank tests compared to the corresponding strain. *, p < 0.05; **, p < 0.01; ***, p < 0.001; NS, not significant, p > 0.05. **e**, IC50 of NTD-targeting NAbs against emerging Omicron subvariants and BA.2 mutants with single NTD substitution.

### Accurate inference of RBD evolution hotspots

Moreover, we wonder if the real-world evolutionary trends of SARS-CoV-2 RBD could be rationalized and even predicted by aggregating this large DMS dataset containing mAbs from various immune histories. Using the mAbs elicited from WT vaccinees or convalescents weighted by IC50 against the D614G strain, we identified mutation hotspots including K417N/T, K444-G446, N450, L452R, and especially E484K (Extended Data Fig. 8a). Most of these residues were mutated in previous VOCs, such as K417N/E484K in Beta, K417T/E484K in Gamma, L452R in Delta, and G446S/E484A in Omicron BA.1, confirming our estimation and inference. Evidence of the emergence of BA.2.75 and BA.5 could also be found using WT, BA.1, and BA.2-elicited mAbs with IC50 against BA.2, where peaks on 444-446, 452, 460, and 486 could be identified (Extended Data Fig. 8c). To better investigate the evolution trends of BA.2.75 and BA.5, the two major lineages circulating currently, we then included antibodies elicited by various immune background, including WT/BA.1/BA.2/BA.5 convalescents, which we believe is the best way to represent the current heterogeneous global humoral immunity (Fig. 4b and Extended Data Fig. 8d). For BA.2.75, the most significant sites are R346T/S, K356T, N417Y/H/I/T, K444E/Q/N/T/M, V445D/G/A, N450T/D/K/S, L452R, I468N, A484P, F486S/V, and F490S/Y. We noticed that these identified residues, even specific mutations, highly overlapped with recent mutation hotspots of BA.2.75 (Fig. 1). Two exceptions are A484 and I468N. E484 is a featured residue of Group C antibodies and could be covered by L452 and F490 (Extended Data Fig. 5c). I468N mutation is also highly associated with K356 mutations, and its function could be covered by K356T (Extended Data Fig. 7a-b). Due to stronger antibody evasion, the preference spectrum of BA.5 is much more concentrated compared to BA.2.75, but the remaining sites are highly overlapped and complementary with BA.2.75. The most striking residues are R346, K444-G446, and N450, followed by K356, N417, L455, N460, and A484. As expected, L452R/F486V does not stand out in BA.5 preference spectrum, while N460K harbored by BA.2.75 appears. These sites and mutations are also popular in emerging BA.4/5 subvariants, proving that our RBD evolution inference system works accurately.

### NAb evasion mechanism of convergent mutations

It is important to examine where this convergent evolution would lead. Based on the observed and predicted convergent hotpots on RBD of BA.2.75 and BA.5, we wonder if we could construct the convergent variants in advance and investigate to what extent they will evade the humoral immune response. To do this, we must first evaluate the antibody evasion mechanism and impact on hACE2-binding capability of the convergent mutations and their combinations. Thus, we selected a panel of 178 NAbs from 8 epitope groups that could potently neutralize BA.2 and determined their neutralizing activity against constructed mutants carrying single or multiple convergent mutations (Fig. 4c, Extended Data Fig. 9a). Most of these sites were selected since we have observed at least 5 independent emergences in distinct lineages of BA.2 and BA.5 that exhibited a growth advantage. NAbs from F1-F3 epitope groups were not included since they are either completely escaped by BA.2 or too rare in Omicron-infected convalescents (Fig. 3f). As expected, R493Q and N417T are not major contributors to antibody evasion, but R493Q significantly benefits ACE2 binding. V445A and K444N caused slightly, and F486S/V caused significantly reduced ACE2-binding capability, consistent with the measurement of emerging subvariants (Fig. 2a, 4c and Extended Data Fig. 1a). The neutralization of NAbs in each group is generally in line with DMS profiles. Most group A NAbs are sensitive to N460K and L455S, and BA.5+N460K escapes the majority of NAbs in group A due to the combination of F486V and N460K (Extended Data Fig. 5a). All NAbs in group B are escaped by F486S/V, and Group C NAbs are heavily escaped by F490S and are strongly affected by L452R and F486S/V (Extended Data Fig. 5b-c). A part of group C NAbs is also slightly affected by K444N/T, S494P and N450D. G446S affects a part of the D1/D2 NAbs, as previously reported ^19^. D1/D2 NAbs are more susceptible to K444N/T, V445A and N450D, and some D1 NAbs could also be escaped by L452R, F490S, and S494P (Extended Data Fig. 6a-b). E1 is mainly affected by R346T, D339H and K356T (Extended Data Fig. 6c). E2.1 and E2.2 exhibit similar properties, evaded by K356T, R346T and L452R (Extended Data Fig. 7a-b). E3 antibodies seem not significantly affected by any of the constructed mutants, as expected (Extended Data Fig. 7c), but they generally exhibit very low neutralization (Extended Data Fig. 7d). BA.5+R346T escapes most antibodies in D1, E1, and E2.1/E2.2, and an additional K444N further escapes most mAbs in D2, demonstrating the feasibility and effectiveness of combining convergent mutations to achieve further evasion. Importantly, adding six mutations to BA.5 could achieve the evasion of the vast majority of RBD NAbs, while exhibiting high hACE2-binding capability, despite the reduction caused by K444N/T and F486V. BQ.1.1, XBB and CH.1.1 could also escape the majority of RBD-targeting NAbs. Together, these findings indicate the feasibility of generating a heavy-antibody-escaping mutant harboring accumulated convergent escape mutations while maintaining sufficient hACE2-binding capability (Fig. 4c and Extended Data Fig. 9a).

Although the proportion of Omicron-specific mAbs is low due to immune imprinting, it is still necessary to evaluate their neutralization potency and breadth, especially against the convergent mutants. We tested the neutralizing activity of a panel of Omicron-specific RBD-targeting mAbs against D614G, BA.1, BA.2, BA.5, BA.2.75, BA.2.75.2, BR.1, BR.2, CA.1, BQ.1.1 and XBB. These mAbs were isolated from convalescent plasma one month after Omicron breakthrough infection (Fig. 4d). They could bind RBD of the corresponding exposed Omicron variant but do not cross-bind WT RBD, as confirmed by ELISA. We found these mAbs could effectively neutralize against the exposed strain, as expected, but exhibited poor neutralizing breadth, which means their potency would be largely impaired by other Omicron subvariants, consistent with our previous discovery ^2^. Notably, BQ.1.1 and XBB could escape most of these Omicron-specific NAbs. Thus, these Omicron-specific antibodies would not effectively expand the breadth of the neutralizing antibody repertoire of Omicron breakthrough infection when facing convergent variants. Further affinity maturation may improve the breadth, but additional experiment is needed.

We then evaluated the potency of NTD-targeting NAbs against BA.2, BA.4/5, BA.2.75 and their sublineages and constructed mutants with selected NTD mutations using a panel of 14 NTD-targeting NAbs, as it is reported that NTD-targeting antibodies are abundant in plasma from BA.2 breakthrough infection and contribute to cross-reactivity ^39^. Most selected mutations are from recently designated Omicron subvariants, except for R237T, which was near V83A, designed to escape mAbs targeting an epitope reported recently ^19^. None of the NTD-targeting NAbs exhibit strong neutralizing potency, and the IC50 values are all over 0.2 μg/mL ^40,41^ (Fig. 4e and Extended Data Fig. 9b). We found the tested BA.2-effective NTD-targeting NAbs could be separated into two clusters, named group α and δ in our previous report, respectively ^19^ (Extended Data Fig. 9c). NAbs in group α target the well-known antigenic supersite on NTD ^42^, which is sensitive to K147E and W152R harbored by BA.2.75*, and Y144del harbored by BJ.1/XBB; while the other group δ is affected by V83A (XBB) and R237T. The other three NTD mutations harbored by BA.2.75, F157L, I210V and G257S did not significantly affect the tested mAbs, consistent with previous sera neutralization data ^17^. Two of the NTD mutations harbored by BJ.1 or XBB, Y144del and V83A, each escapes a cluster of them and together would enable XBB to exhibit extremely strong capability of escaping NTD-targeting NAbs. Notably, XBB.1 escaped all NTD-targeting NAbs tested here.

### Simulation of convergent variant evolution

Based on the above results, we designed multiple VSV-based pseudoviruses that gradually gain convergent mutations that could induce RBD/NTD-targeting NAb resistance (Fig. 5a). The constructed final mutant contains 11 additional mutations on the NTD and RBD compared to BA.5, or 9 mutations compared to BA.2.75. The neutralizing activities of Omicron-effective NAb drugs were first evaluated. As expected, the majority of existing effective NAb drugs, except SA55, are escaped by these mutants (Fig. 5b). Similarly, we also determined the ACE2-binding capability of these mutants by neutralization assays using hACE2 (Fig. 5c). Although some of the designed pseudoviruses, especially those with K444N and F486V, exhibit reduced activity to hACE2 compared to original BA.2.75 or BA.5 variants, their binding affinities are still higher than that of D614G (Fig. 2b). Importantly, our designed pseudoviruses could largely evade the plasma of vaccinees and convalescents after BA.1 breakthrough infection, BA.2 breakthrough infection, and even BA.5 breakthrough infection (Fig. 5d-g). Among the derivative mutants of BA.2.75, L452R, K444M, R346T, and F486V contribute mainly to the significant reduction in neutralization (Fig. 5d-g). Adding more NTD mutations does not contribute to stronger evasion in BA.2.75-based mutants, but we observed a significant reduction in NT50 of BA.2/BA.5 convalescents against BA.5-based mutants with K147E+W152R, suggesting BA.2/BA.5 convalescent plasma contains a large proportion of NTD-targeting antibodies ^39^. As the NTD of BA.1 differs from that of BA.2 and BA.5, we did not observe significant effects of NTD mutations on the efficacy of BA.1 convalescent plasma. Plasma neutralization titers of most vaccinees and convalescents decreased to the lower detection limit against BA.2.75 with 5 extra RBD mutations L452R, K444M, R346T, F486V, and K356T. The same applies to vaccinees or BA.1 convalescents against BA.5 with 4 extra RBD mutations K444N, R346T, N460K, and K356T. The plasma from BA.2/BA.5 convalescents can tolerate more mutations based on BA.5, and extra NTD mutations such as K147E and W152R are needed to completely eliminate their neutralization. Together, we demonstrate that as few as five additional mutations on BA.5 or BA.2.75 could completely evade most plasma samples, including those from BA.5 breakthrough infection, while maintaining high hACE2-binding capability. Similar efforts have been made in a recent report despite different construction strategies ^43^. The constructed evasive mutants, such as BA.2.75-S5/6/7/8 and BA.5-S7/8, could serve to examine the effectiveness of broad-spectrum vaccines and NAbs in advance.

**Fig. 5.**
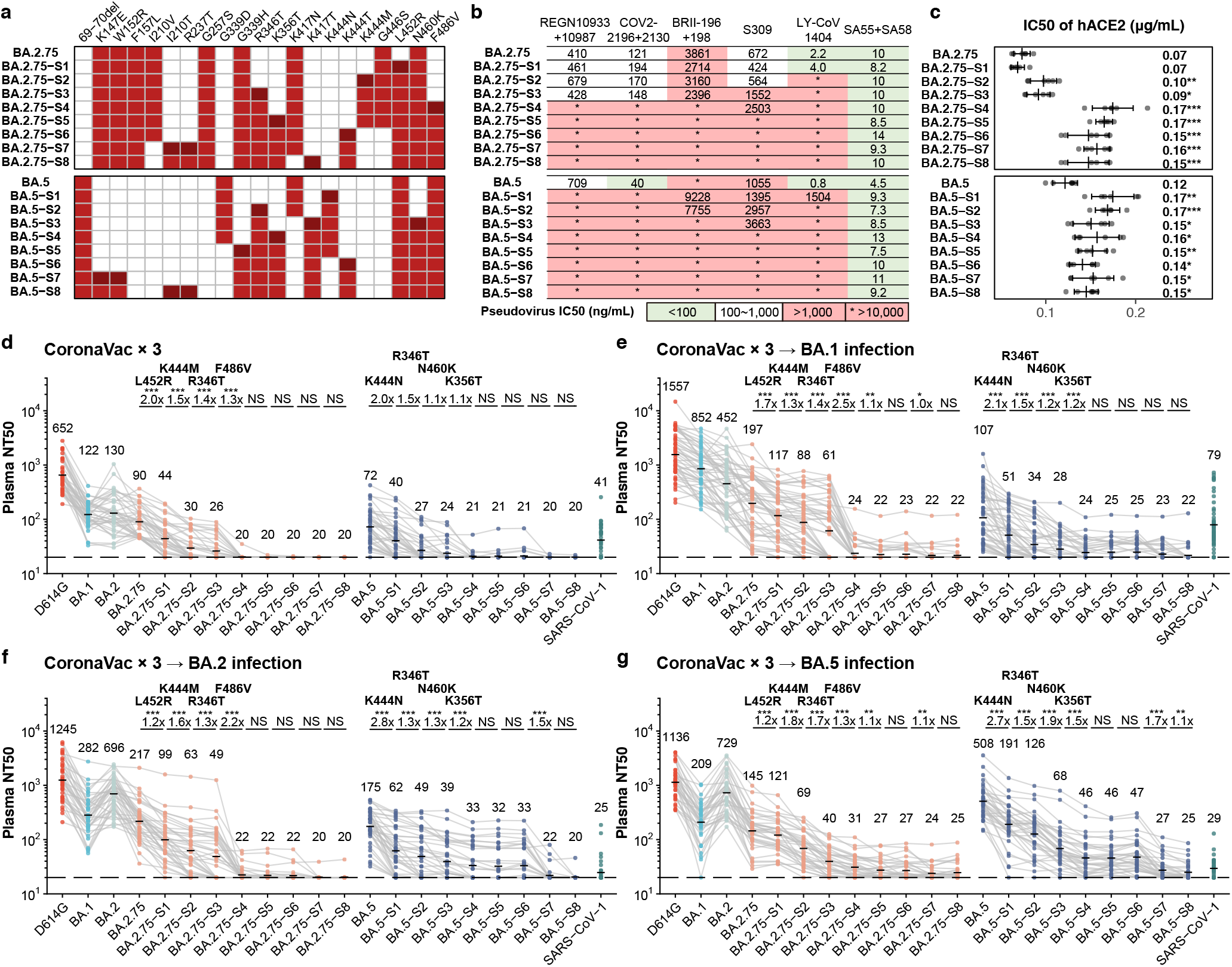
Accumulation of convergent escape mutations leads to complete loss of plasma neutralization. **a**, Mutations of multiple designed mutants that harbor key convergent escape mutations based on BA.2.75 and BA.5. **b**, IC50 of therapeutic mAbs and cocktails against pseudoviruses of designed mutants. green, IC50 ≤ 100ng/mL; white, 100ng/mL < IC50 < 1,000ng/mL; red, IC50 ≥ 1,000ng/mL; *, IC50 ≥ 10,000ng/mL. **c**, IC50 of hACE2 against the designed mutants. Error bars indicate mean±s.d. P-values were calculated using two-tailed Student’s t-test, compared to BA.2.75 and BA.5 respectively for BA.2.75 and BA.5-derived mutants. *, p < 0.05; **, p < 0.01; ***, p < 0.001. No label on variants with p > 0.05. **d-g**, Pseudovirus neutralizing titers against SARS-CoV-2 D614G, Omicron subvariants and designed mutants of plasma from vaccinated or convalescent individuals from breakthrough infection. **d**, Individuals who received 3 doses of CoronaVac (n = 40). **e**, Convalescents infected with BA.1 after receiving 3 doses of CoronaVac (n = 50). **f**, Convalescents infected with BA.2 after receiving 3 doses of CoronaVac (n = 39). **g**, Convalescents infected with BA.5 after receiving 3 doses of CoronaVac (n = 36). Key additional mutations harbored by each designed mutant are annotated above the points. The geometric mean titers are labeled. P-values are determined using two-tailed Wilcoxon signed-rank tests of paired samples. *, p < 0.05; **, p < 0.01; ***, p < 0.001; NS, not significant, p > 0.05. Statistical tests are determined using two-tailed Wilcoxon signed-rank tests.

## Discussion

In this work, we showed that convergent RBD evolution can cause severe immune evasion and could be rationalized by integrating DMS profiles. Given the existence of immune imprinting, the humoral immune repertoire is not effectively diversified by infection with new Omicron variants, while the immune pressure on the RBD becomes increasingly concentrated and promotes convergent evolution, posing a great challenge to current vaccines and antibody drugs.

Although this study only examines inactivated vaccines, immune imprinting is also observed in those receiving mRNA vaccines ^44,45^. In fact, mRNA-vaccinated individuals displayed an even higher proportion of cross-reactive memory B cells, probably because the overall humoral immune response induced by mRNA vaccines is stronger than that induced by inactivated vaccines ^45^. Also, recent studies on mRNA vaccinees who receive a BA.5 booster or BA.5 breakthrough infection displayed similar neutralization reduction trend against BA.2.75.2, BQ.1 and BQ.1.1, suggesting high consistency of neutralization data among vaccine types ^46,47^.

As the antibodies undergo affinity maturation, their SHM rate would increase ^45^. This may lead to a higher proportion of variant-specific antibodies, enhanced binding affinity, and increased neutralization breadth, which could potentially resist the convergent mutations carried by variants like XBB and BQ.1.1 ^48^. However, the effect of affinity maturation may be counteracted by waning immunity ^45,49^. The affinity-matured memory B cells would require a second booster or reinfection to be effectively deployed.

We also observed that plasma from convalescents with BA.5 breakthrough infection exhibited higher neutralization against BA.5-derived variants like BQ.1 and BQ.1.1, suggesting that BA.5-based boosters are beneficial to protection against convergent variants of BA.5 sublineages. However, this may be mainly driven by the enrichment of NTD-targeting antibodies after BA.5 breakthrough infection, which was also reported in BA.2 convalescents ^39^. Significant mutations on NTD, such as Y144del in XBB and BQ.1.18, and mutations of many BA.2.75 sublineages, would cause severe reduction in BA.5 breakthrough infection plasma neutralization titers. Therefore, the effectiveness of BA.5-based boosters against the convergent mutants carrying critical NTD mutations should be closely monitored.

Notably, the antibody evasion capability of many variants, such as BQ.1.1, CA.1, BQ.1.18, XBB, and CH.1.1, have already reached or even exceeded SARS-CoV-1, indicating extensive antigenic drift (Fig. 5d-g). Indeed, by constructing an antigenic map of the tested SARS-CoV-2/SARS-CoV-1 variants using the plasma NT50 data, we found that the antigenicity distances of SARS-CoV-2 ancestral strain to CA.1, CH.1.1, XBB and BQ.1.1 are already comparable to that of SARS-CoV-1 (Extended Data Fig. 10a-b). Given that there are ~50 different amino acids between SARS-CoV-1 and SARS-CoV-2 RBD, but only 21 mutations on BQ.1.1 RBD compared to the ancestral strain, these results indicate that the global pandemic indeed has greatly promoted the efficiency of the virus to evolve immune escape mutations.

Finally, our prediction demonstrated a remarkable consistency with real-world observations. Some variants close to the predicted and constructed variants have already emerged while we performed the experiments, validating our prediction model. For example, BQ.1.1 is highly similar to BA.5-S3, and CH.1.1 to BA.2.75-S4/S6 (Fig. 4c). The whole pipeline for constructing pseudoviruses carrying predicted mutations could be safely conducted in biosafety level 2 (BSL-2) laboratories, and does not involve any infectious pandemic virus. If we had this prediction model at the beginning of the pandemic, the development of NAb drugs and vaccines might not be so frustrated against the continuously emerging SARS-CoV-2 variants. Broad-spectrum SARS-CoV-2 vaccines and NAb drugs development should be of high priority, and the constructed convergent mutants could serve to examine their effectiveness in advance.

## Methods

### Isolation of peripheral blood mononuclear cells and plasma

Samples from vaccinees and individuals who had recovered from BA.1, BA.2, or BA.5 infection were obtained under study protocols approved by Beijing Ditan Hospital, Capital Medical University (Ethics committee archiving No. LL-2021-024-02) and the Tianjin Municipal Health Commission, and the Ethics Committee of Tianjin First Central Hospital (Ethics committee archiving No. 2022N045KY). All donors provided written informed consent for the collection of information, the use of blood and blood components, and the publication of data generated from this study. Whole blood samples were diluted 1:1 with PBS+2%FBS (Gibco) and subjected to Ficoll (Cytiva) gradient centrifugation. Plasma was collected from the upper layer. Cells were collected at the interface and further prepared by centrifugation, red blood cell lysis (Invitrogen eBioscience) and washing steps. The date of vaccination, hospitalization and sampling can be found in Supplementary Table 1.

### BCR sequencing, analysis, and antibody production

CD19+ B cells were isolated from PBMCs with EasySep Human CD19 Positive Selection Kit II (STEMCELL, 17854). Every 10^6^ B cells in 100 μl solution were stained with 3 μl FITC antihuman CD20 antibody (BioLegend, 302304, clone: 2H7), 3.5 μl Brilliant Violet 421 antihuman CD27 antibody (BioLegend, 302824, clone: O323), 2 μl PE/Cyanine7 anti-human IgM antibody (BioLegend, 314532, clone: MHM-88), 2 μl PE/Cyanine7 anti-human IgD antibody (BioLegend, 348210, clone: IA6-2), 0.13 μg biotinylated SARS-CoV-2 BA.2 RBD protein (customized from Sino Biological) or 0.13 μg biotinylated SARS-CoV-2 BA.5 RBD protein (customized from Sino Biological) conjugated with PE-streptavidin (BioLegend, 405204) and APC-streptavidin (BioLegend, 405207), 0.13 μg SARS-CoV-2 WT biotinylated RBD protein (Sino Biological, 40592-V27H-B) conjugated with Brilliant Violet 605 Streptavidin (BioLegend, 405229). Cells are also labeled with biotinylated RBD conjugated to DNA-oligo-streptavidin. Omicron RBD (BA.2 or BA.5) were labeled with TotalSeq-C0971 Streptavidin (Biolegend, 405271) and TotalSeq-C0972 Streptavidin (Biolegend, 405273); WT RBD were labled with TotalSeq-C0973 Streptavidin (Biolegend, 405275) and TotalSeq-C0974 Streptavidin (Biolegend, 405277). Cells were washed twice after 30 minutes of incubation on ice. 7-AAD (Invitrogen, 00-6993-50) was used to label dead cells. 7-AAD-CD20+CD27+IgM-IgD-SARS-CoV-2 BA.2 RBD+ or BA.5 RBD+ cells were sorted with a MoFlo Astrios EQ Cell Sorter. FACS data were analyzed using FlowJo v10.8 (BD Biosciences).

Sorted B cells were resuspended in the appropriate volume and then processed with Chromium Next GEM Single Cell V(D)J Reagent Kits v1.1 following the manufacturer’s user guide (10x Genomics, CG000208). Gel beads-in-emulsion (GEMs) were obtained with a 10X Chromium controller. GEMs were subjected to reverse transcription and purification. Reverse transcription products were subject to preamplification and purification with SPRIselect Reagent Kit (Beckman Coulter, B23318). BCR sequences (paired V(D)J) were enriched with 10X BCR primers. After library preparation, libraries were sequenced with the Illumina sequencing platform.

10X Genomics V(D)J sequencing data were assembled as BCR contigs and aligned using Cell Ranger (v6.1.1) pipeline according to the GRCh38 BCR reference. Only the productive contigs and the B cells with one heavy chain and one light chain were kept for quality control. The germline V(D)J gene identification and annotation were performed by IgBlast (v1.17.1)^51^. Somatic hypermutation sites in the antibody variable domain were detected using Change-O toolkit (v1.2.0)^52^.

Antibody heavy and light chain genes were optimized for human cell expression and synthesized by GenScript. VH and VL were inserted separately into plasmids (pCMV3-CH, pCMV3-CL or pCMV3-CK) through infusion (Vazyme, C112). Plasmids encoding the heavy chain and light chain of antibodies were co-transfected by polyethylenimine-transfection to Expi293F™ cell (ThermoFisher, A14527). Cells were cultured at 36.5°C, 5% CO2, 175 rpm for 6-10 days. Supernatants containing mAbs were collected, and the supernatants were further purified with protein A magnetic beads (Genscript, L00695).

### High-throughput deep mutation scanning

High-throughput DMS platform has been described previously^1,2^. Briefly, deep mutation scanning libraries were constructed by mutagenesis PCR based on the Wuhan-Hu-1 RBD sequence (GenBank: MN908947, residues N331-T531). A unique 26-nucleotide (N26) barcode was appended to each RBD variant in mutant libraries by PCR, and the correspondence between the N26 barcode and mutations in RBD variants was acquired by PacBio sequencing. RBD mutant libraries were first transformed in the EBY100 strain of *Saccharomyces cerevisiae* and then enriched for properly folded ACE2 binders, which were used for subsequent mutation escape profiling. The above ACE2 binders were grown in SG-CAA media (2% w/v d-galactose, 0.1% w/v dextrose (d-glucose), 0.67% w/v yeast nitrogen base, 0.5% w/v casamino acids (-ade, -ura, -trp), 100 mM phosphate buffer, pH 6.0) at room temperature for 16-18h with agitation. Then these yeast cells were washed twice and proceeded to three rounds of magnetic beads-based selection. Obtained yeast cells after sequential sorting were recovered overnight in SD-CAA media (2% w/v dextrose (d-glucose), 0.67% w/v yeast nitrogen base, 0.5% w/v casamino acids (-ade, -ura, -trp), 70 mM citrate buffer, pH 4.5). Pre- and post-sort yeast populations were submitted to plasmid extraction by 96 Well Plate Yeast Plasmid Preps Kit (Coolaber, PE053). N26 barcode sequences were amplified with the extracted plasmid templates, and PCR products were purified and submitted to Illumina Nextseq 550 sequencing.

### Antibody clustering and embedding based on DMS profiles

Data analysis of DMS was performed as described in previous reports ^1,2^. In brief, the detected barcode sequences of both the antibody-screened and reference library were aligned to the barcode-variant lookup table generated using dms_variants (v0.8.9). The escape scores of each variant X in the library were defined as F×(n_X,ab_ / N_ab_) / (n_X,ref_ / N_ref_), where F is a scale factor to normalize the scores to the 0-1 range, while n and N are the number of detected barcodes for variant X and total barcodes in post-selected (ab) or reference (ref) samples, respectively. The escape scores of each mutation were calculated by fitting an epistasis model as described previously ^37,53^.

Epitope groups of new antibodies not included in our previous report are determined by k-nearest neighbors (KNN)-based classification. In brief, site escape scores of each antibody are first normalized and considered as a distribution across RBD residues, and only residues whose standard derivation is among the highest 50% of all residues are retained for further analysis. Then the dissimilarity or distance of two antibodies is defined by the Jessen-Shannon divergence of the normalized escape scores. Pair-wise dissimilarities of all antibodies in the dataset are calculated using the scipy package (scipy.spatial.distance.jensenshannon, v1.7.0). For each antibody, 15 nearest neighbors whose epitope groups have been determined by unsupervised clustering in our previous paper are identified and simply voted to determine the group of the selected antibody. To project the dataset onto a 2D space for visualization, we performed MDS to represent each antibody in a 32-dimensional space, and then t-SNE to get the 2D representation, using sklearn.manifold.MDS and sklearn.manifold.TSNE (v0.24.2).

### Calculation of the estimated preference of RBD mutations

Four different weights are included in the calculation, including the weight for ACE2-binding affinity, RBD expression, codon constraint, and neutralizing activity. Impact on ACE2-binding affinity and RBD expression of each mutation based on WT, BA.1 and BA.2 are obtained from public DMS results. And for BA.5 (BA.2+L452R+F486V+R493Q) and BA.2.75 (BA.2+D339H+G446S+N460K+R493Q), BA.2 results are used except for these mutated residues, whose scores for each mutant are subtracted by the score for the mutation in BA.5 or BA.2.75. As the reported values are log fold changes, the weight is simply defined by the exponential of reported values, i.e., exp [*S*_bind_] or exp[*S*_expr_], respectively. For codon constraint, the weight is 1.0 for mutants that could be accessed by one nucleotide mutation, and 0.0 for others. We used the following RBD nucleotide sequences for determination of accessible mutants, WT/D614G (Wuhan-Hu-1 reference genome), BA.1 (EPI_ISL_10000028), BA.2 (EPI_ISL_10000005), BA.4/5 (EPI_ISL_11207535), BA.2.75 (EPI_ISL_13302209). For neutralizing activity, the weight is -log_10_(IC50). The IC50 values (μg/mL), which are smaller than 0.0005 or larger than 1.0 are considered as 0.0005 or 1.0, respectively. The raw escape scores for each antibody are first normalized by the max score among all mutants, and the final weighted score for each antibody and each mutation is the production of the normalized scores and four corresponding weights. The final mutation-specific weighted score is the summation of scores of all antibodies in the designated antibody set, and then normalized again to make it a value between 0 and 1. Logo plots for visualization of escape maps were generated by the Python package logomaker (v0.8).

### Pseudovirus neutralization assay

The Spike gene (GenBank: MN908947) was mammalian condon-optimized and inserted into the pcDNA3.1 vector. Site-directed mutagenesis PCR was performed as described previously^54^. The sequence of mutants is shown in Supplementary Table 3. Pseudotyped viruses were generated by transfection 293T cells (ATCC, CRL-3216) with pcDNA3.1-Spike with Lipofectamine 3000 (Invitrogen). The cells were subsequently infected with G*ΔG-VSV (Kerafast) that packages expression cassettes for firefly luciferase instead of VSV-G in the VSV genome. The cell supernatants were discarded after 6-8h harvest and replaced with complete culture media. The cell was cultured for one day, and then the cell supernatant containing pseudotyped virus was harvested, filtered (0.45-μm pore size, Millipore), aliquoted, and stored at −80 °C. Viruses of multiple variants were diluted to the same number of copies before use.

mAbs or plasma was serially diluted and incubated with the pseudotyped virus in 96-well plates for 1 h at 37°C. Trypsin-treated Huh-7 cells (Japanese Collection of Research Bioresources, 0403) were added to the plate. The cells were cultured for 20-28 h in 5% CO_2_, 37°C incubators. The supernatants were removed and left 100 μL in each well, and 100 μL luciferase substrate (Perkinelmer, 6066769) was added and incubated in the dark for 2 min. The cell lysate was removed, and the chemiluminescence signals were collected by PerkinElmer Ensight. Each experiment was repeated at least twice.

Inhibitory efficiencies of hACE2 against the pseudoviruses were determined with the same procedure, using hACE2-Fc dimer (Sino Biological, 10108-H02H), and each experiment was conducted in five biological replicates.

Dulbecco’s modified Eagle medium (DMEM, high glucose; HyClone) with 100 U/mL of penicillin-streptomycin solution (Gibco), 20 mM N-2-hydroxyethylpiperazine-N-2-ethane sulfonic acid (HEPES, Gibco) and 10% fetal bovine serum (FBS, Gibco) were used in cell culture. Trypsin-EDTA (0.25%, Gibco) was used to detach cells before seeding to the plate.

### Enzyme-linked immunosorbent assay

WT/BA.2/BA.1 RBD or spikes in PBS was pre-coated onto ELISA plates at 4 °C overnight and were washed and blocked. 1 μg ml^-1^ purified antibodies or serially diluted antibodies were added and incubated at room temperature for 20 min. Peroxidase-conjugated AffiniPure Goat Anti-Human IgG (H+L) (JACKSON, 109-035-003) was added to plates and incubated at room temperature for 15 min. Tetramethylbenzidine (TMB) (Solarbio, 54827-17-7) was added and incubated for 10 mins, and then the reaction was terminated with 2 M H2SO4. Absorbance was measured at 450 nm using a microplate reader (PerkinElmer, HH3400).

## Acknowledgments

We thank J. Bloom for his gift of the yeast SARS-CoV-2 RBD libraries. We thank all volunteers for providing the blood samples. We thank all the scientists around the globe for performing SARS-CoV-2 sequencing and surveillance analysis. This project is financially supported by the Ministry of Science and Technology of China and Changping Laboratory (2021A0201; 2021D0102), and National Natural Science Foundation of China (32222030).

## Author contributions

Y.C. designed the study. X.S.X supervised the study. Y.C, F.J., A.Y, Q.G. and X.S.X. wrote the manuscript with inputs from all authors. A.Y., W.S., R.A., Yao W., and X.N. performed B-cell sorting, single-cell VDJ sequencing, and antibody sequence analyses. J.W. (BIOPIC), H.S., and F.J. performed and analyzed the DMS data. Y.Y. and Youchun W. constructed the pseudotyped virus. N.Z., P.W., L.Y., T.X. and F.S. performed the pseudotyped virus neutralization assays. W.S. and Y.C. analyzed the neutralization data. X.H., Y.X., X.C., Z.S. and R.J. recruited the SARS-CoV-2 vaccinees and convalescents. J.W. (Changping Laboratory), L.Y. and F.S. performed the antibody expression.

## Data and Code Availability

Processed mutation escape scores and custom scripts to analyze the data can be downloaded at https://github.com/jianfcpku/convergent_RBD_evolution. Sequences and neutralization of the antibodies are included in Supplementary Table 2. Any other raw data of specific antibodies are available from the corresponding authors upon request. We used vdj_GRCh38_alts_ensembl-5.0.0 as the reference of V(D)J alignment, which can be obtained from https://support.10xgenomics.com/single-cell-vdj/software/downloads/latest. IMGT/DomainGapAlign is based on the built-in latest IMGT antibody database, and we let the “Species” parameter as “Homo sapiens” while keeping the others as default.

## Extended Data Figures

**Extended Data Fig. 1.**
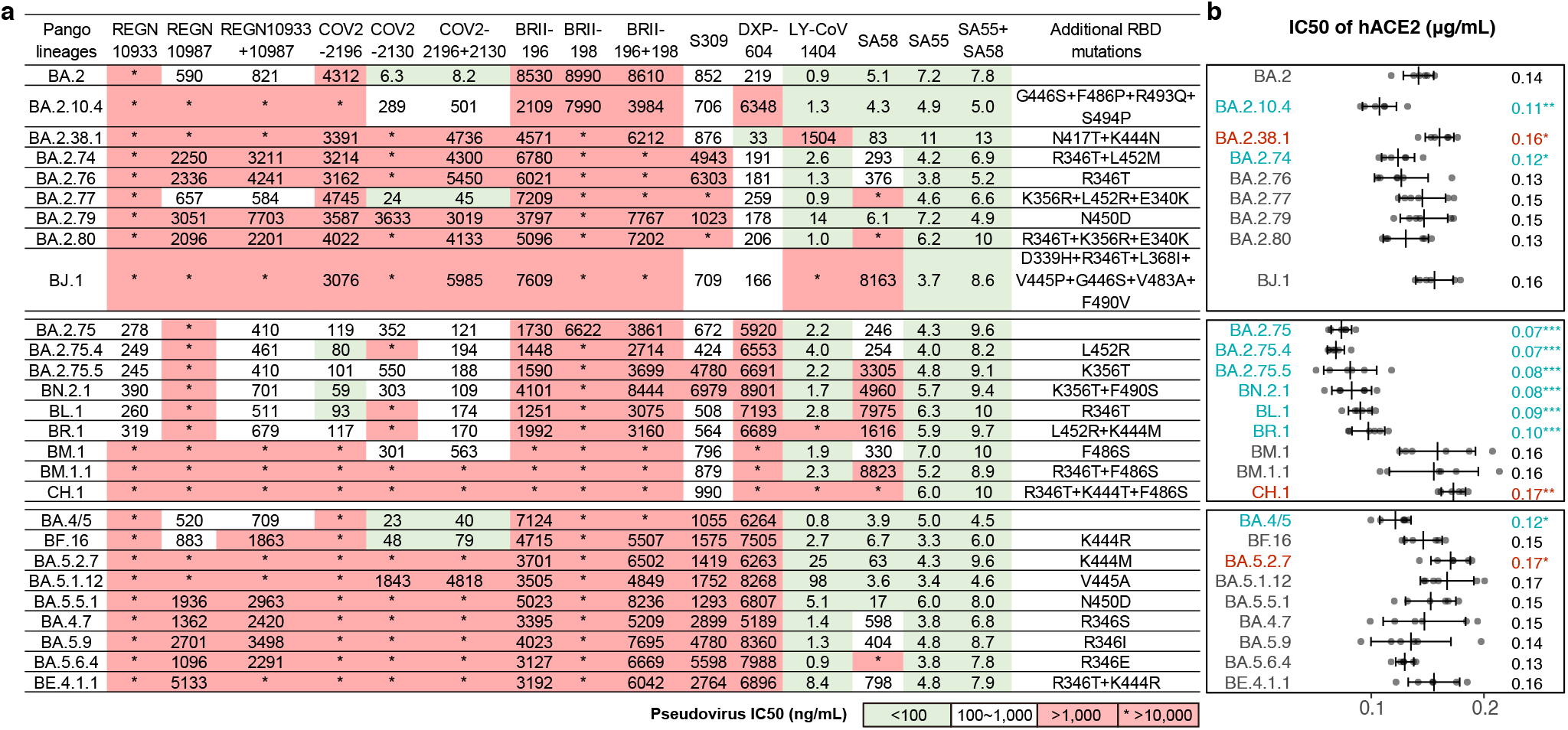
Antibody drug evasion and hACE2 binding capability of convergent Omicron variants. **a**, IC50 of therapeutic NAbs against pseudoviruses of additional emerging SARS-CoV-2 Omicron subvariants. **b**, Relative hACE2-binding capability measured by IC50 of hACE2 against pseudoviruses. Error bars indicate mean±s.d. P-values were calculated using a twotailed Student’s *t*-test. *, p < 0.05; **, p < 0.01; ***, p < 0.001. No label on variants with p > 0.05. Variants with significantly stronger binding are colored blue, while those with weaker binding are colored red.

**Extended Data Fig. 2.**
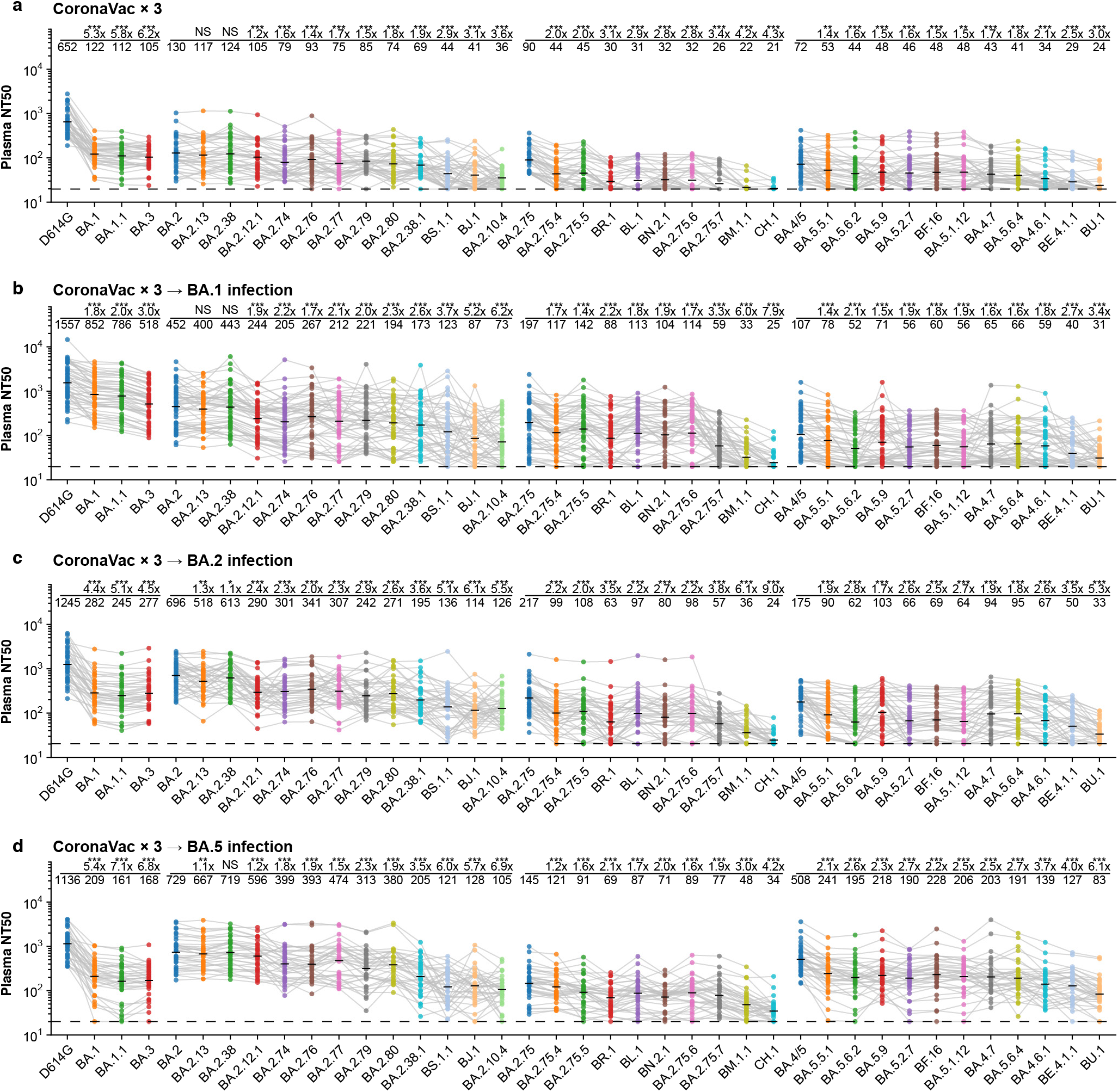
Plasma neutralization evasion of convergent Omicron variants. **a-d**, NT50 against SARS-CoV-2 previous variants of concern and additional Omicron subvariants of plasma from vaccinated individuals or convalescents of breakthrough infection. Plasma samples, statistical methods and meaning of labels are the same as in Fig. 2.

**Extended Data Fig. 3.**
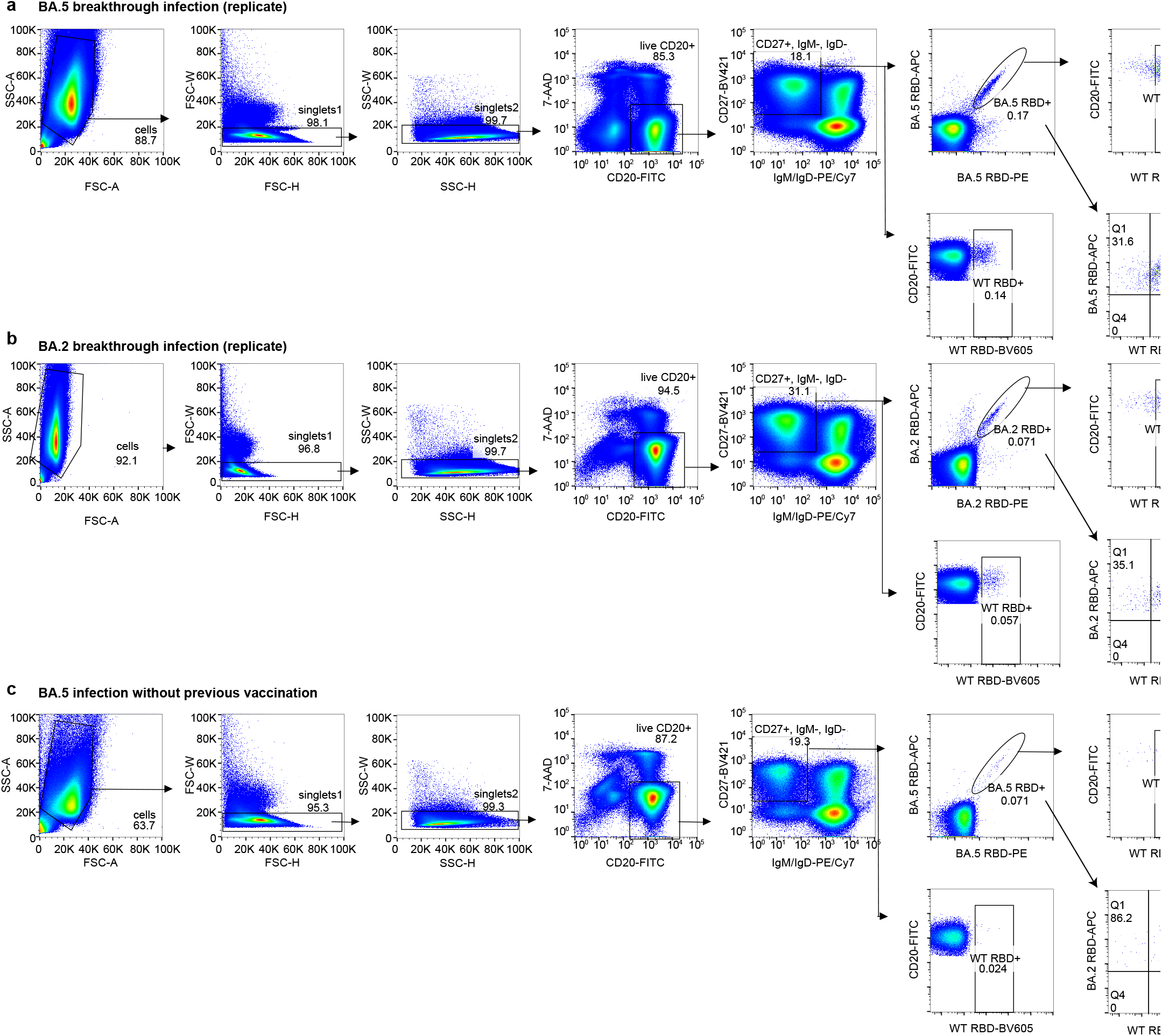
FACS gating strategy for isolating mAbs from BA.2 and BA.5 convalescents. **a**, FACS gating strategy of antigen-specific B cells from individuals who recovered from BA.5 breakthrough infection. Data from an independent experiment compared to Fig. 3a are shown here. **b**, FACS gating strategy of antigen-specific B cells from individuals who recovered from BA.2 breakthrough infection. Data from an independent experiment compared to Fig. 3b are shown here. **c**, FACS gating strategy of antigen-specific B cells from individuals who recovered from BA.5 infection.

**Extended Data Fig. 4.**
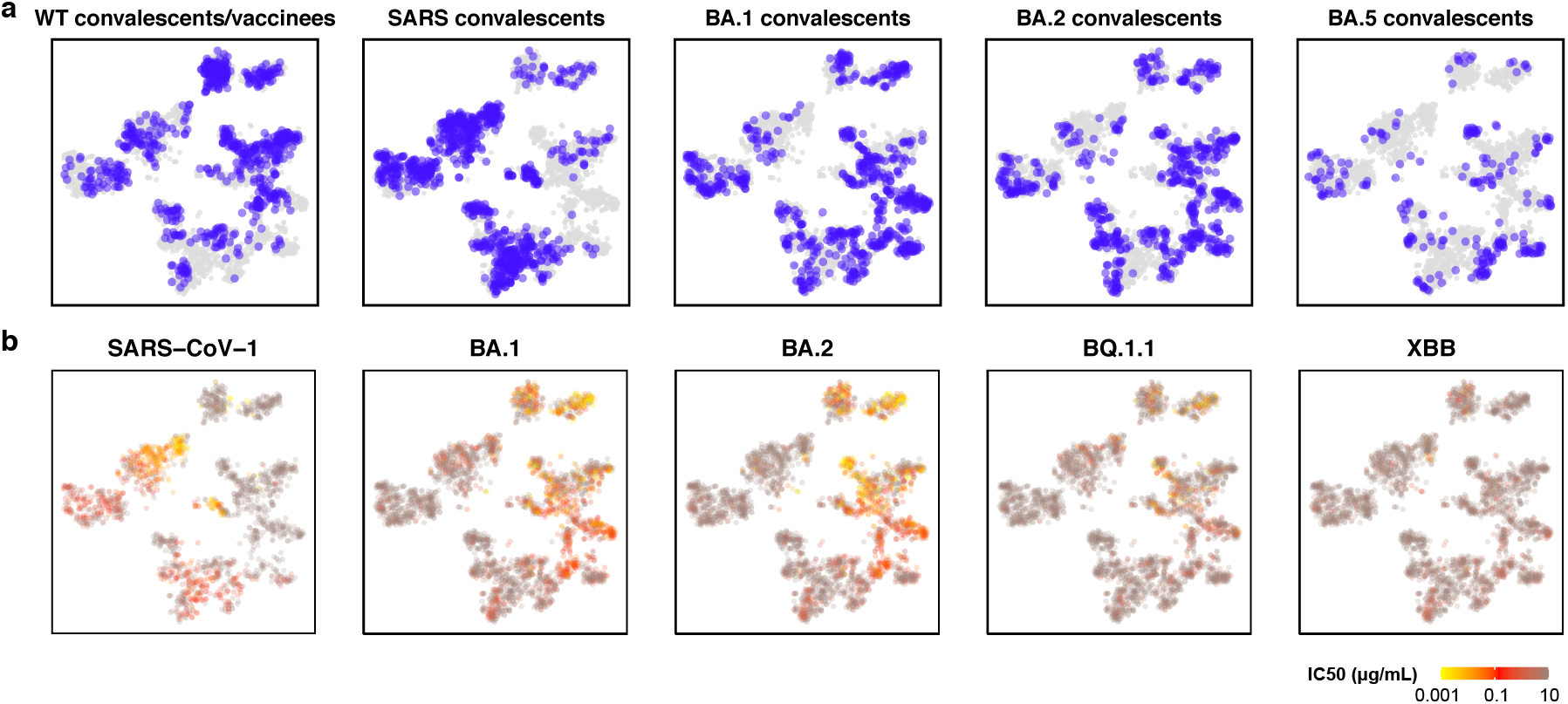
Distribution of antibody sources and neutralizing activities on the DMS landscape. **a**, Sources of the 3051 mAbs involved in this study projected on the t-SNE of DMS profiles. **b**, IC50 against SARS-CoV-1 (N=1870 determined), Omicron BA.1 (N=3031), BA.2 (N=3046), BQ.1.1 (N=3051), and XBB (N=3033) of these mAbs projected on the embedding.

**Extended Data Fig. 5.**
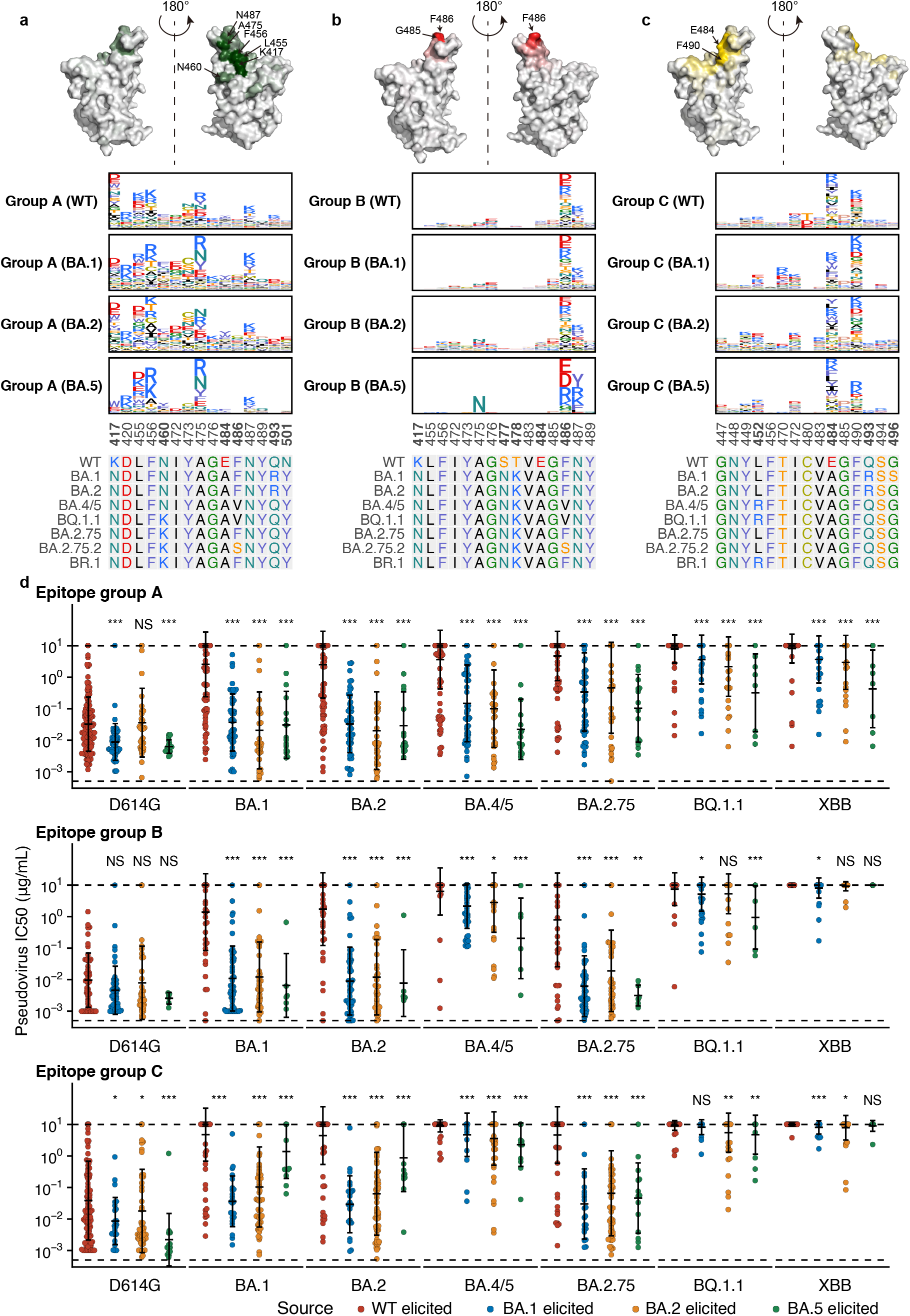
Escape hotspots and neutralization of mAbs in epitope groups A, B and C. **a-c,** Average escape scores from DMS of epitope groups A (**a**), B (**b**), C (**c**) and each RBD residue. Scores are projected onto the structure of SARS-CoV-2 RBD (PDB: 6M0J). Average escape maps that indicate the score of each mutation from DMS on escape hotspots of antibodies, grouped by their sources, in epitope groups A (**a**), B (**b**) and C (**c**), and corresponding sequence alignment of SARS-CoV-2 WT and Omicron RBDs are also shown. The height of each amino acid in the escape maps represents its mutation escape score. Mutated sites in Omicron variants are marked in bold. **d**, Pseudovirus-neutralizing IC50 of antibodies in group A, B, and C, from wildtype convalescents or vaccinees (WT-elicited, n=133, 50, 106 for A-C, respectively), BA.1 convalescents (BA.1-elicited, n=51, 49, 24), BA.2 convalescents (BA.2-elicited, n=34, 36, 56) and BA.5 convalescents (BA.5-elicited, n=16, 6, 14). The geometric mean IC50s are labeled, and error bars indicate the geometric standard deviation. P-values are calculated using two-tailed Wilcoxon signed-rank tests. *, p < 0.05; **, p < 0.01; ***, p < 0.001; NS, not significant, p > 0.05.

**Extended Data Fig. 6.**
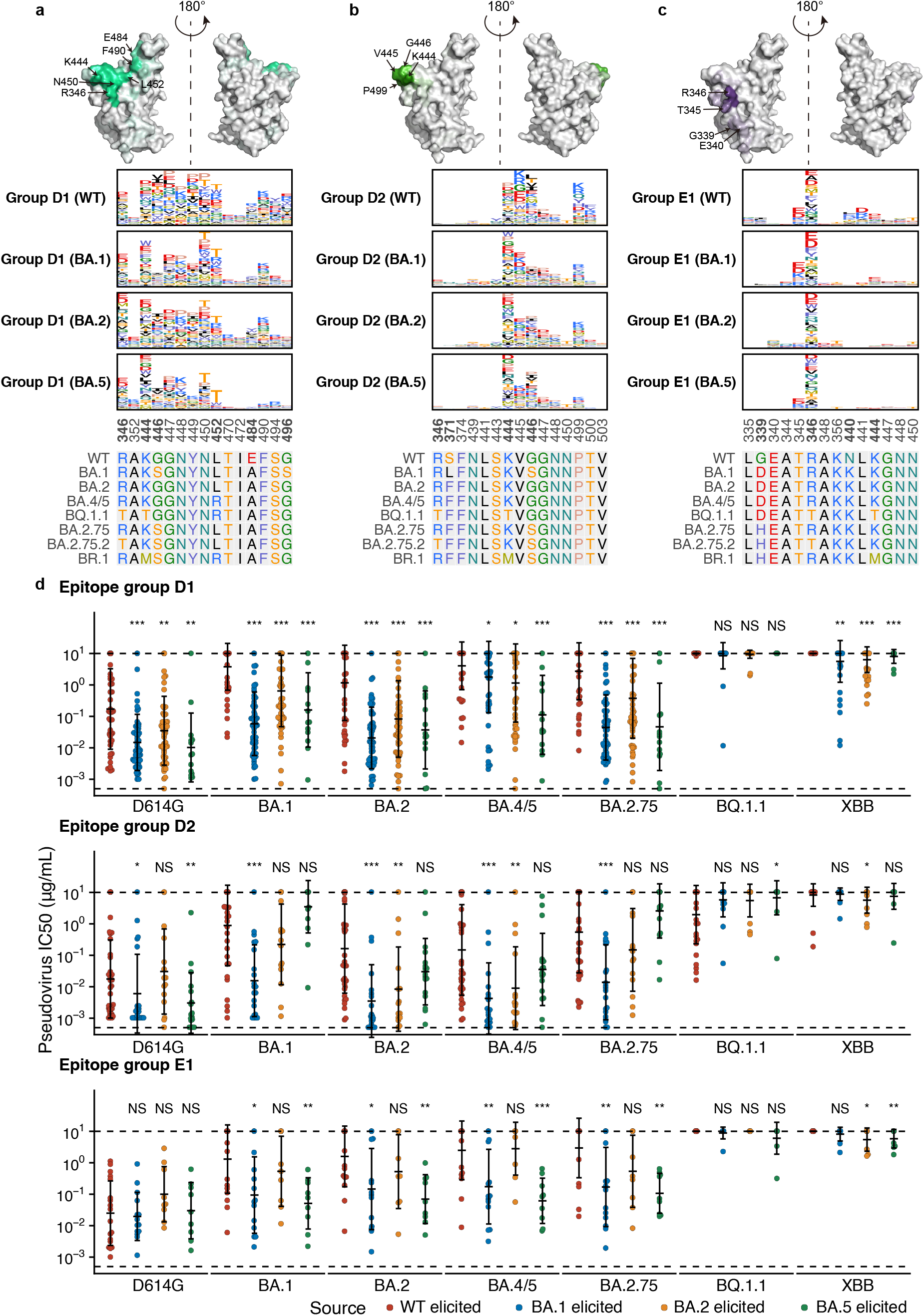
Escape hotspots and neutralization of mAbs in epitope group D and E1. **a-c,** Average escape scores from DMS of epitope groups D1 (**a**), D2 (**b**), E1 (**c**) and each RBD residue. **d**, Pseudovirus-neutralizing IC50 of antibodies in group D1, D2, and E1 For WT-elicited mAbs (n=49, 37, 19 for D1, D2 and E1, respectively), BA.1 convalescents (n=59, 21, 14 for D1, D2 and E1, respectively), BA.2 convalescents (n=56, 15, 9 for D1, D2 and E1, respectively), and BA.5 convalescents (n=14, 17, 9 for D1, D2 and E1, respectively).

**Extended Data Fig. 7.**
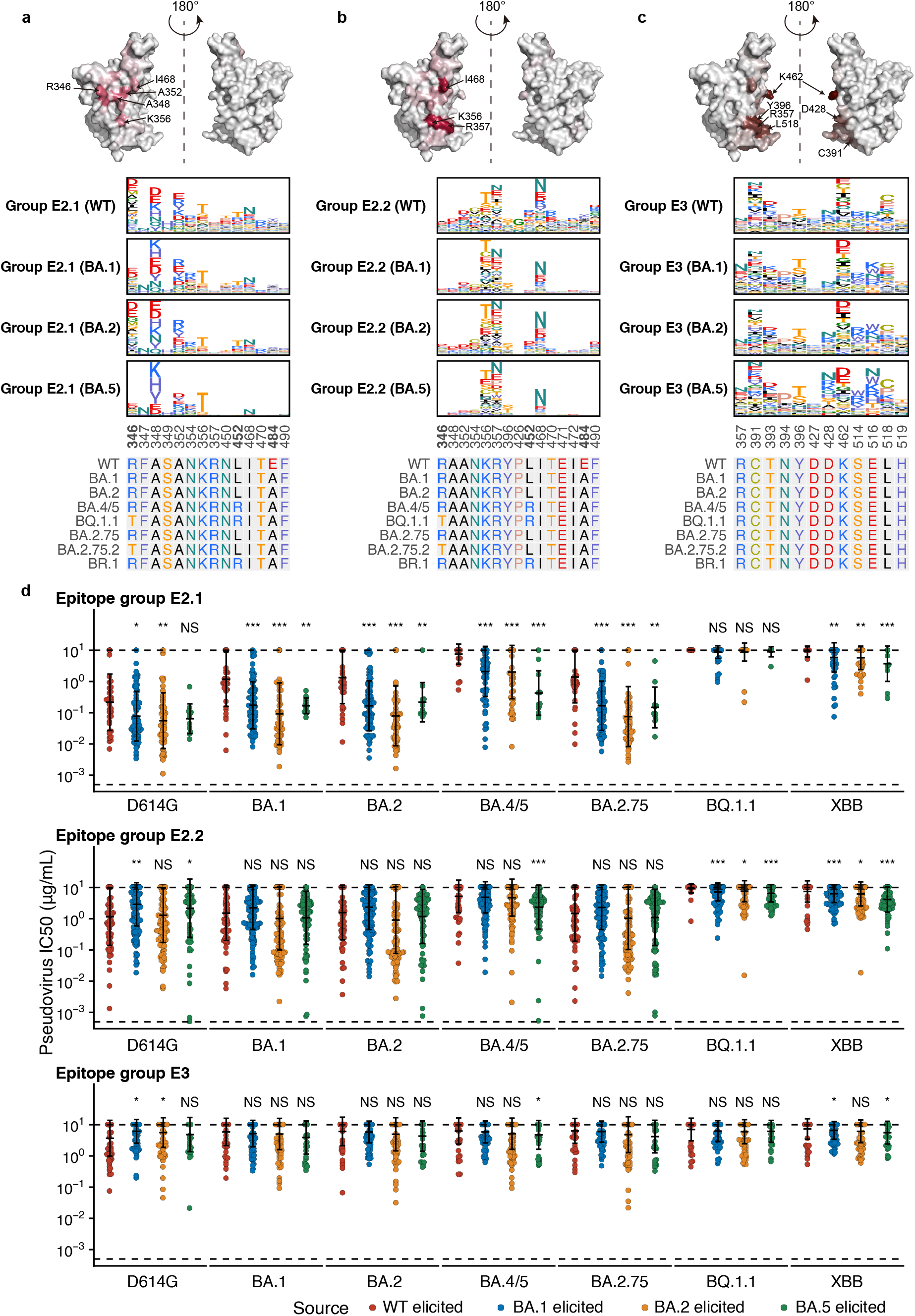
Escape hotspots and neutralization of mAbs in epitope group E2 and E3. **a-c,** Average escape scores from DMS of epitope groups E2.1 (**a**), E2.2 (**b**), E3 (**c**) and each RBD residue. **d**, Pseudovirus-neutralizing IC50 of antibodies in group E2.1, E2.2, and E3 for WT-elicited mAbs (n=49, 37, 19 for E2.1, E2.2, and E3, respectively), BA.1 convalescents (n=59, 21, 14 for E2.1, E2.2, and E3, respectively), BA.2 convalescents (n=56, 15, 9 for E2.1, E2.2, and E3, respectively), and BA.5 convalescents (n=14, 17, 9 for E2.1, E2.2, and E3, respectively).

**Extended Data Fig. 8.**
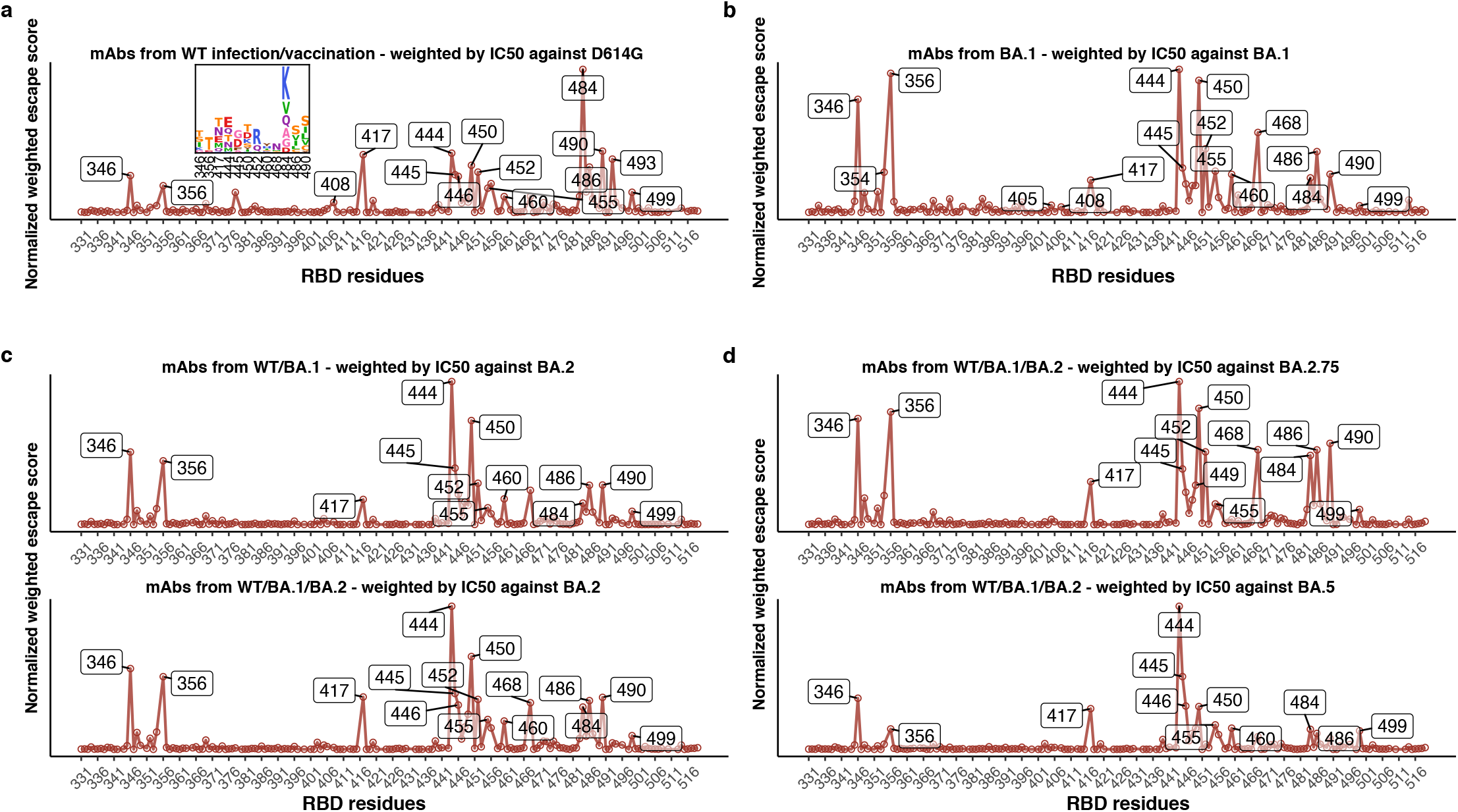
Predicted escape hotspots of SARS-CoV-2 variants. **a**, Normalized average escape scores weighted by IC50 against D614G using DMS profiles of mAbs from ancestral strain infection or vaccination with a logo plot showing specific mutations on important residues. **b**, Normalized average escape scores of mAbs from BA. 1 breakthrough infection, weighted by IC50 against BA.1. **c**, Normalized average escape scores of mAbs from ancestral strain infection or vaccination and BA.1 breakthrough infection, weighted by IC50 against BA.2. **d**, WT/BA.1/BA.2-elicited mAbs with IC50 against BA.2.75 and BA.5, similar to Fig. 4b.

**Extended Data Fig. 9.**
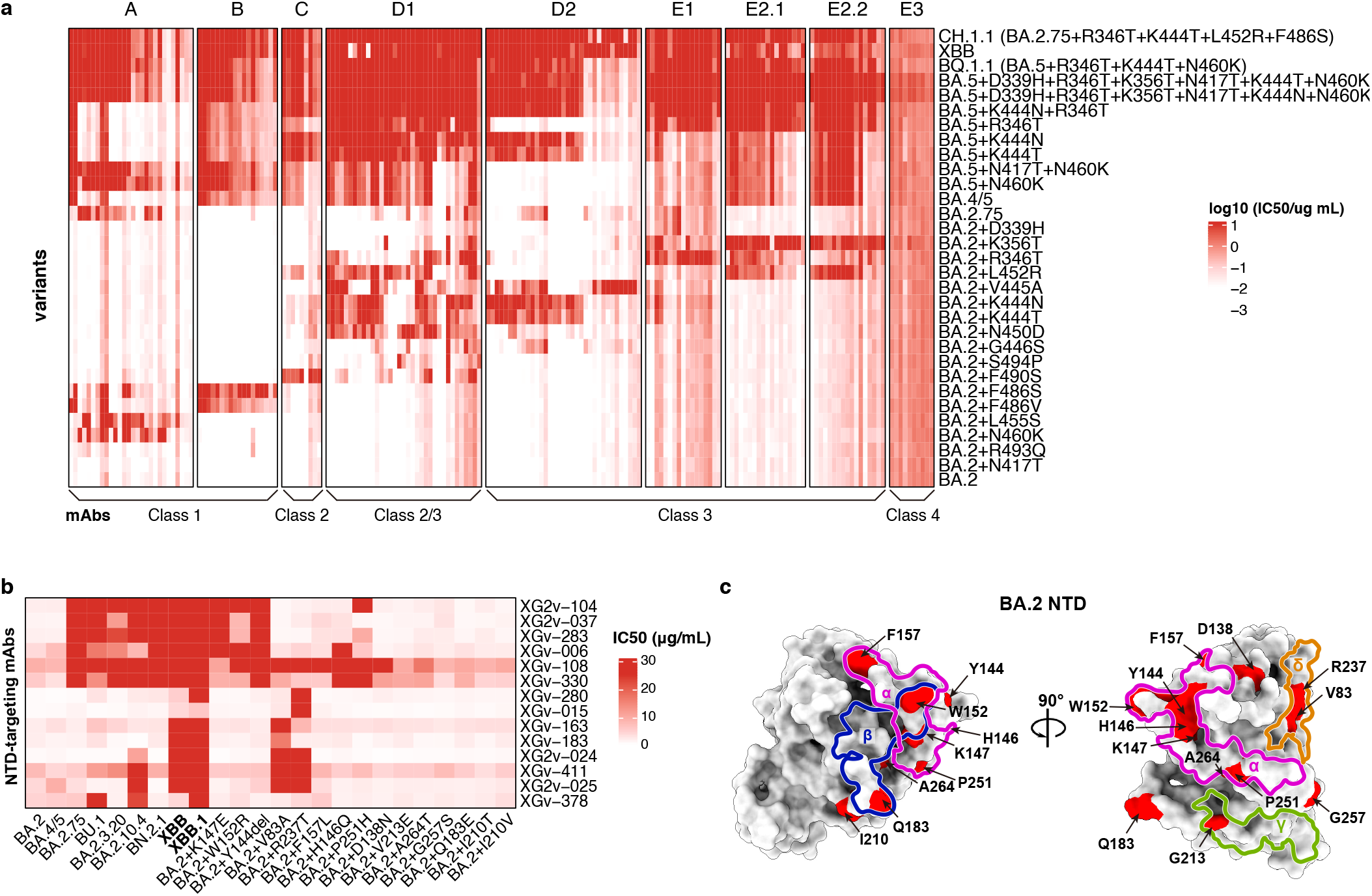
IC50 heatmaps of representative mAbs against constructed Omicron variants. **a**, Color shades indicate IC50 of antibodies (columns) against constructed Omicron BA.2 or BA.5 subvariants (rows) carrying mutations on the epitope of each group. The order of mAbs is the same as in Figure 4c. **b**, IC50 of NTD-targeting antibodies against SARS-CoV-2 variants, which is related to Fig. 4e. **c**, Epitope groups and escape hotspots on BA.2 NTD.

**Extended Data Fig. 10.**
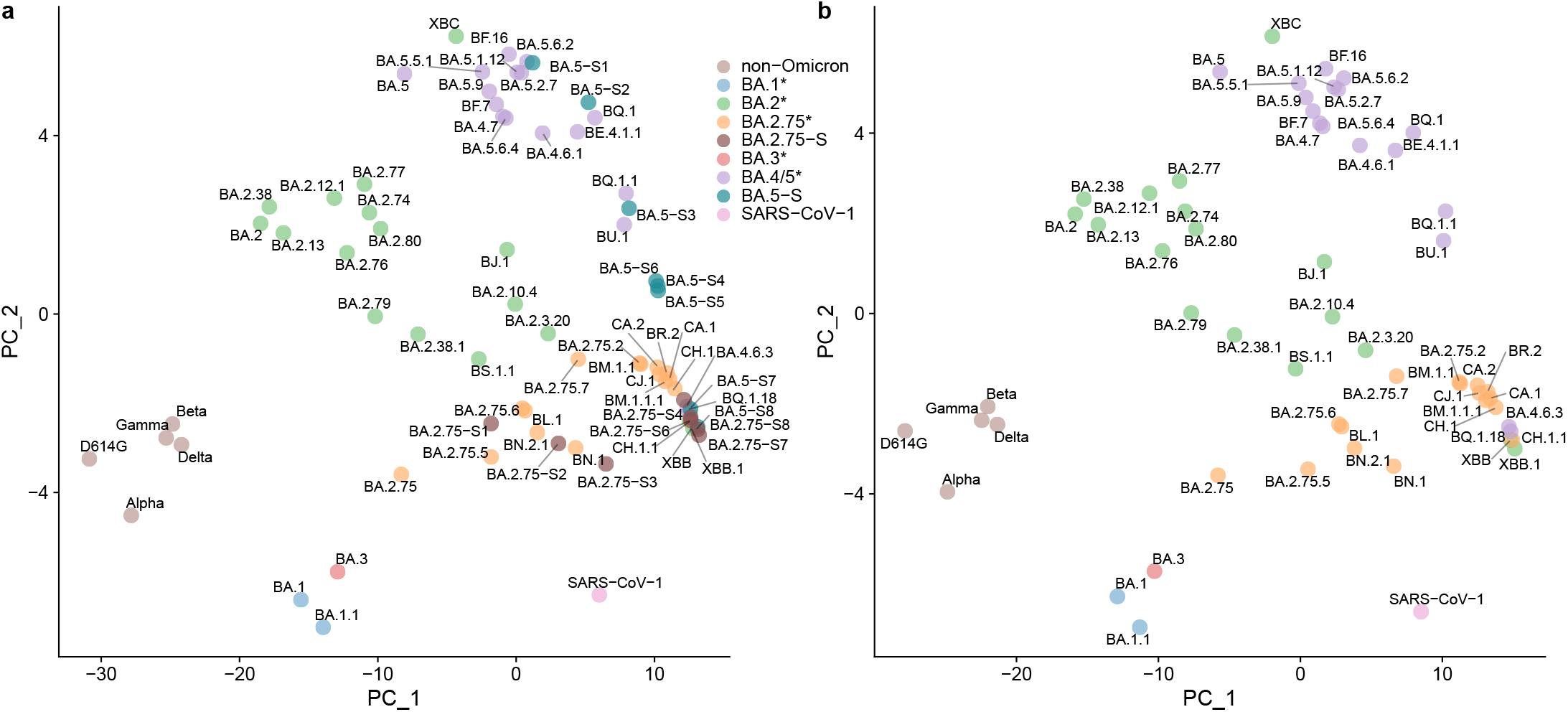
Antigenic map of current SARS-CoV-2 variants. **a**, Antigenic map of SARS-CoV-2 variants constructed using plasma neutralization data by principal component analysis (PCA). **b**, Antigenic map of SARS-CoV-2 variants with constructed Omicron subvariants removed.

## References

1 Cao, Y. et al. Omicron escapes the majority of existing SARS-CoV-2 neutralizing antibodies. Nature 602, 657–663, doi:10.1038/s41586-021-04385-3 (2022).

2 Cao, Y. et al. BA.2.12.1, BA.4 and BA.5 escape antibodies elicited by Omicron infection. Nature 608, 593–602, doi:10.1038/s41586-022-04980-y (2022).

3 Wang, Q. et al. Antibody evasion by SARS-CoV-2 Omicron subvariants BA.2.12.1, BA.4 and BA.5. Nature 608, 603–608, doi:10.1038/s41586-022-05053-w (2022).

4 Tuekprakhon, A. et al. Antibody escape of SARS-CoV-2 Omicron BA.4 and BA.5 from vaccine and BA.1 serum. Cell 185, 2422–2433 e2413, doi:10.1016/j.cell.2022.06.005 (2022).

5 Uraki, R. et al. Characterization and antiviral susceptibility of SARS-CoV-2 Omicron/BA.2. Nature, doi:10.1038/s41586-022-04856-1 (2022).

6 Liu, L. et al. Striking antibody evasion manifested by the Omicron variant of SARS-CoV-2. Nature 602, 676–681, doi:10.1038/s41586-021-04388-0 (2022).

7 Cele, S. et al. Omicron extensively but incompletely escapes Pfizer BNT162b2 neutralization. Nature 602, 654–656, doi:10.1038/s41586-021-04387-1 (2022).

8 Cameroni, E. et al. Broadly neutralizing antibodies overcome SARS-CoV-2 Omicron antigenic shift. Nature 602, 664–670, doi:10.1038/s41586-021-04386-2 (2022).

9 Yamasoba, D. et al. Virological characteristics of the SARS-CoV-2 Omicron BA.2 spike. Cell 185, 2103–2115 e2119, doi:10.1016/j.cell.2022.04.035 (2022).

10 Nutalai, R. et al. Potent cross-reactive antibodies following Omicron breakthrough in vaccinees. Cell 185, 2116–2131 e2118, doi:10.1016/j.cell.2022.05.014 (2022).

11 Dejnirattisai, W. et al. SARS-CoV-2 Omicron-B.1.1.529 leads to widespread escape from neutralizing antibody responses. Cell 185, 467–484 e415, doi:10.1016/j.cell.2021.12.046 (2022).

12 Cui, Z. et al. Structural and functional characterizations of infectivity and immune evasion of SARS-CoV-2 Omicron. Cell 185, 860–871 e813, doi:10.1016/j.cell.2022.01.019 (2022).

13 Wang, K. et al. Memory B cell repertoire from triple vaccinees against diverse SARS-CoV-2 variants. Nature 603, 919–925, doi:10.1038/s41586-022-04466-x (2022).

14 Kimura, I. et al. Virological characteristics of the SARS-CoV-2 Omicron BA.2 subvariants including BA.4 and BA.5. Cell, doi:10.1016/j.cell.2022.09.018 (2022).

15 Shu, Y. & McCauley, J. GISAID: Global initiative on sharing all influenza data - from vision to reality. Eurosurveillance 22, 30494, doi:doi:https://doi.org/10.2807/1560-7917.ES.2017.22.13.30494 (2017).

16 Rambaut, A. et al. A dynamic nomenclature proposal for SARS-CoV-2 lineages to assist genomic epidemiology. Nature Microbiology 5, 1403–1407, doi:10.1038/s41564-020-0770-5 (2020).

17 Wang, Q. et al. Antigenic characterization of the SARS-CoV-2 Omicron subvariant BA.2.75. Cell Host & Microbe, doi:https://doi.org/10.1016/j.chom.2022.09.002 (2022).

18 Sheward, D. J. et al. Evasion of neutralising antibodies by omicron sublineage BA.2.75. Lancet Infect Dis, doi:10.1016/S1473-3099(22)00524-2 (2022).

19 Cao, Y. et al. Characterizations of enhanced infectivity and antibody evasion of Omicron BA.2.75. bioRxiv, 2022.2007.2018.500332, doi:10.1101/2022.07.18.500332 (2022).

20 Jian, F. et al. Further humoral immunity evasion of emerging SARS-CoV-2 BA.4 and BA.5 subvariants. bioRxiv, 2022.2008.2009.503384, doi:10.1101/2022.08.09.503384 (2022).

21 Saito, A. et al. Virological characteristics of the SARS-CoV-2 Omicron BA.2.75. bioRxiv, 2022.2008.2007.503115, doi:10.1101/2022.08.07.503115 (2022).

22 Chen, C. et al. CoV-Spectrum: Analysis of Globally Shared SARS-CoV-2 Data to Identify and Characterize New Variants. Bioinformatics 38, 1735–1737, doi:10.1093/bioinformatics/btab856 (2021).

23 Greaney, A. J. et al. Comprehensive mapping of mutations in the SARS-CoV-2 receptor-binding domain that affect recognition by polyclonal human plasma antibodies. Cell Host Microbe 29, 463–476 e466, doi:10.1016/j.chom.2021.02.003 (2021).

24 Greaney, A. J. et al. Complete Mapping of Mutations to the SARS-CoV-2 Spike Receptor-Binding Domain that Escape Antibody Recognition. Cell Host Microbe 29, 44–57 e49, doi:10.1016/j.chom.2020.11.007 (2021).

25 Greaney, A. J. et al. Mapping mutations to the SARS-CoV-2 RBD that escape binding by different classes of antibodies. Nature Communications 12, 4196, doi:10.1038/s41467-021-24435-8 (2021).

26 Zost, S. J. et al. Potently neutralizing and protective human antibodies against SARS-CoV-2. Nature 584, 443–449, doi:10.1038/s41586-020-2548-6 (2020).

27 Westendorf, K. et al. LY-CoV1404 (bebtelovimab) potently neutralizes SARS-CoV-2 variants. Cell Rep 39, 110812, doi:10.1016/j.celrep.2022.110812 (2022).

28 Cao, Y. et al. Rational identification of potent and broad sarbecovirus-neutralizing antibody cocktails from SARS convalescents. bioRxiv, 2022.2008.2003.499114, doi:10.1101/2022.08.03.499114 (2022).

29 Loo, Y. M. et al. The SARS-CoV-2 monoclonal antibody combination, AZD7442, is protective in nonhuman primates and has an extended half-life in humans. Sci Transl Med 14, eabl8124, doi:10.1126/scitranslmed.abl8124 (2022).

30 Starr, T. N. et al. Deep mutational scans for ACE2 binding, RBD expression, and antibody escape in the SARS-CoV-2 Omicron BA.1 and BA.2 receptor-binding domains. bioRxiv, 2022.2009.2020.508745, doi:10.1101/2022.09.20.508745 (2022).

31 Gao, Q. et al. Development of an inactivated vaccine candidate for SARS-CoV-2. Science 369, 77–81, doi:10.1126/science.abc1932 (2020).

32 Quandt, J. et al. Omicron BA.1 breakthrough infection drives cross-variant neutralization and memory B cell formation against conserved epitopes. Sci Immunol 7, eabq2427, doi:10.1126/sciimmunol.abq2427 (2022).

33 Khan, K. et al. Omicron BA.4/BA.5 escape neutralizing immunity elicited by BA.1 infection. Nature Communications 13, 4686, doi:10.1038/s41467-022-32396-9 (2022).

34 Park, Y. J. et al. Imprinted antibody responses against SARS-CoV-2 Omicron sublineages. bioRxiv, 2022.2005.2008.491108, doi:10.1101/2022.05.08.491108 (2022).

35 Reynolds, C. J. et al. Immune boosting by B.1.1.529 (Omicron) depends on previous SARS-CoV-2 exposure. Science 377, eabq1841, doi:10.1126/science.abq1841 (2022).

36 Greaney, A. J., Starr, T N. & Bloom, J. D. An antibody-escape estimator for mutations to the SARS-CoV-2 receptor-binding domain. Virus Evol 8, veac021, doi:10.1093/ve/veac021 (2022).

37 Starr, T N. et al. Deep Mutational Scanning of SARS-CoV-2 Receptor Binding Domain Reveals Constraints on Folding and ACE2 Binding. Cell 182, 1295–1310 e1220, doi:10.1016/j.cell.2020.08.012 (2020).

38 Starr, T N. et al. Shifting mutational constraints in the SARS-CoV-2 receptor-binding domain during viral evolution. Science 377, 420–424, doi:10.1126/science.abo7896 (2022).

39 Muik, A. et al. Omicron BA.2 breakthrough infection enhances cross-neutralization of BA.2.12.1 and BA.4/BA.5. Science Immunology 0, eade2283, doi:10.1126/sciimmunol.ade2283 (2022).

40 Wang, Z. et al. Analysis of memory B cells identifies conserved neutralizing epitopes on the N-terminal domain of variant SARS-Cov-2 spike proteins. Immunity 55, 998–1012 e1018, doi:10.1016/j.immuni.2022.04.003 (2022).

41 Chi, X. et al. A neutralizing human antibody binds to the N-terminal domain of the Spike protein of SARS-CoV-2. Science 369, 650–655, doi:doi:10.1126/science.abc6952 (2020).

42 McCallum, M. et al. N-terminal domain antigenic mapping reveals a site of vulnerability for SARS-CoV-2. Cell 184, 2332–2347 e2316, doi:10.1016/j.cell.2021.03.028 (2021).

43 Witte, L. et al. Epistasis lowers the genetic barrier to SARS-CoV-2 neutralizing antibody escape. bioRxiv, 2022.2008.2017.504313, doi:10.1101/2022.08.17.504313 (2022).

44 Alsoussi, W. B. et al. SARS-CoV-2 Omicron boosting induces de novo B cell response in humans. bioRxiv, 2022.2009.2022.509040, doi:10.1101/2022.09.22.509040 (2022).

45 Kaku, C. I. et al. Evolution of antibody immunity following Omicron BA.1 breakthrough infection. bioRxiv, 2022.2009.2021.508922, doi:10.1101/2022.09.21.508922 (2022).

46 Qu, P. et al. Distinct Neutralizing Antibody Escape of SARS-CoV-2 Omicron Subvariants BQ.1, BQ.1.1, BA.4.6, BF.7 and BA.2.75.2. bioRxiv, 2022.2010.2019.512891, doi:10.1101/2022.10.19.512891 (2022).

47 Wang, Q. et al. Antibody responses to Omicron BA.4/BA.5 bivalent mRNA vaccine booster shot. bioRxiv, 2022.2010.2022.513349, doi:10.1101/2022.10.22.513349 (2022).

48 Marzi, R. et al. Maturation of SARS-CoV-2 Spike-specific memory B cells drives resilience to viral escape. bioRxiv, 2022.2009.2030.509852, doi:10.1101/2022.09.30.509852 (2022).

49 Levin, E. G. et al. Waning Immune Humoral Response to BNT162b2 Covid-19 Vaccine over 6 Months. New England Journal of Medicine 385, e84, doi:10.1056/NEJMoa2114583 (2021).

50 Barnes, C. O. et al. SARS-CoV-2 neutralizing antibody structures inform therapeutic strategies. Nature 588, 682–687, doi:10.1038/s41586-020-2852-1 (2020).

## Methods references

51 Ye, J., Ma, N., Madden, T L. & Ostell, J. M. IgBLAST: an immunoglobulin variable domain sequence analysis tool. Nucleic Acids Res 41, W34–40, doi:10.1093/nar/gkt382 (2013).

52 Gupta, N. T. et al. Change-O: a toolkit for analyzing large-scale B cell immunoglobulin repertoire sequencing data. Bioinformatics 31, 3356–3358, doi:10.1093/bioinformatics/btv359 (2015).

53 Otwinowski, J., McCandlish, D. M. & Plotkin, J. B. Inferring the shape of global epistasis. Proc Natl Acad Sci U S A 115, E7550–E7558, doi:10.1073/pnas.1804015115 (2018).

54 Nie, J. et al. Quantification of SARS-CoV-2 neutralizing antibody by a pseudotyped virus-based assay. Nat Protoc 15, 3699–3715, doi:10.1038/s41596-020-0394-5 (2020).

